# Selective repurposing of the eukaryotic DNA replication machinery by a plant virus

**DOI:** 10.64898/2026.07.23.740281

**Authors:** Chaonan Shi, Laura Medina-Puche, Delphine M. Pott, Hua Wei, Sarah A. Acatay, Björn Krenz, Linda Hanley-Bowdoin, Rosa Lozano-Durán

**Author notes:** These authors contributed equally to this work (Laura Medina-Puche, Delphine M. Pott).

## Abstract

Eukaryotic DNA viruses that replicate in the nucleus often exploit components of the host DNA replication machinery for genome replication. Geminiviruses, causal agents of devastating crop diseases worldwide, strictly depend on host factors to replicate their circular single-stranded (ss) DNA genomes, with only a single virus-encoded protein, Rep, required for this process. Rep recruits host replication proteins to the viral genome and catalyzes nicking and ligation at the initiation and termination sites of rolling-circle replication. Despite the reliance of geminiviral replication on plant proteins, the composition of the viral replisome remains largely unknown. Here, we use TurboID-based proximity labeling to identify plant proteins in the vicinity of the Rep proteins from the geminiviruses tomato yellow leaf curl virus (TYLCV) and abutilon mosaic virus (AbMV) during infection. Combining virus-induced gene silencing, infection assays, and chromatin immunoprecipitation, we identify host DNA replication-related factors required for viral genome replication and likely components of the viral replisome. Our results indicate that geminiviruses and related eukaryotic ssDNA viruses selectively repurpose components of the eukaryotic replication fork to support rolling-circle replication, and suggest that they follow a leading-strand replication mode while utilizing the lagging-strand DNA polymerase δ. These findings shed light on the molecular mechanism of geminiviral DNA replication and identify potential targets for engineering antiviral resistance in crops.

## INTRODUCTION

Eukaryotic DNA viruses that replicate in the nucleus of infected cells frequently hijack components of the host DNA replication machinery to mediate their own genome replication. This is the case for geminiviruses, plant-infecting viruses that cause devastating crop diseases worldwide. Geminiviruses strictly rely on host factors to replicate their circular single-stranded (ss) DNA genomes, with only a single virus-encoded protein, the Replication-associated protein (Rep), directly involved in this process. Rep reprograms the cell cycle by interfering with the retinoblastoma (RBR in plants, for Retinoblastoma-Related)/E2F pathway, thereby promoting the availability of the necessary molecular machinery. Rep has also been proposed to provide helicase activity (Choudhury *et al*., 2006; Clérot and Bernardi, 2006; George *et al*., 2014; Ruhel *et al*., 2021) to unwind the viral DNA during replication. In addition, Rep recruits host replication-related proteins to the viral genome and catalyses nicking and ligation during the initiation and termination of rolling-circle replication (RCR) (Hanley-Bowdoin *et al*., 2013). Despite this strong dependence on host proteins, the molecular identity of the components of the viral replisome remains largely elusive.

The family *Geminiviridae* is among the largest infecting plants, and some of its members are major pathogens of cash and staple crops. Geminiviruses cause diseases that can lead to severe yield losses, with both economic and humanitarian impacts, posing a serious threat to food security in current and future scenarios (Varma and Malathi, 2003; Rojas *et al*., 2018). Because viral replication is a fundamental step in infection and depends on host molecular machinery, it represents a key vulnerability that that could be exploited for antiviral strategies. Moreover, dissecting how geminiviruses subvert host DNA replication processes can provide fundamental insights into plant cell biology, immunity, and virus–host co-evolution.

Geminiviruses have circular ssDNA genomes ranging from 2.5 to 5.6 kb in size. Upon entry into the host cell, the viral ssDNA is transported to the nucleus, where it is converted into a double-stranded (ds) DNA intermediate with the help of plant DNA polymerase (pol) α (Zhao *et al*., 2019; Wu *et al*., 2021; Bonnamy *et al*., 2023). This dsDNA associates with nucleosomes to form a minichromosome, which serves as a template for both viral gene transcription and RCR (Jeske, 2009; Hanley-Bowdoin *et al*., 2013). RCR requires Rep, which is thought to recruit the necessary host machinery to the viral genome. A second mode of replication described for geminiviruses is recombination-dependent replication (RDR), for which Rep is dispensable, which seems to contribute to viral accumulation in combination with RCR (Preiss and Jeske, 2003). DNA pol δ has been shown to be required for RCR (Wu *et al*., 2021). Although several host DNA replication factors —including proliferating cell nuclear antigen (PCNA), its loader replication factor C (RFC), and the ssDNA-binding replication protein A (RPA)— have been reported to physically interact with geminiviral Rep proteins (Luque *et al*., 2002a; Castillo *et al*., 2003; Bagewadi *et al*., 2004; Singh *et al*., 2007), robust genetic evidence demonstrating their requirement for viral DNA replication remains limited. Consequently, the full repertoire of host proteins required for this process is still incompletely defined.

This replication strategy, in which a virus-encoded Rep protein enables host machinery–mediated RCR, is shared across members of the phylum of eukaryotic Circular Rep-encoding Single-Stranded DNA (CRESS) viruses (*Cressdnaviricota*). At present, this phylum comprises twenty-four families infecting diverse hosts (International Committee for the Taxonomy of Virsues; https://ictv.global/), including four plant-infecting families, *Geminiviridae*, *Nanoviridae*, *Metaxyviridae*, and *Amesuviridae* (Krupovic *et al*., 2020; Gronenborn *et al*., 2021; Da Silva *et al*., 2023). Interestingly, members of the animal-infecting *Circoviridae* family also rely on DNA polymerases α and δ for replication, as later shown for geminiviruses in plants and at least for DNA polymerase δ in their insect vectors (Gassmann *et al*., 1988; He *et al*., 2020; Wu *et al*., 2021). This suggests that the molecular mechanisms underlying CRESS virus replication are likely conserved across families and hosts.

Here, we take advantage of TurboID-based proximity labelling (PL) *in planta* to investigate the composition of the geminiviral replisome. Specifically, we compare the proxiomes of two Rep proteins encoded by tomato yellow leaf curl virus (TYLCV) and abutilon mosaic virus (AbMV) in the context of infection. The requirement of candidate host proteins common to both datasets, as well as additional DNA replication-related factors, was assessed through reverse genetics using tobacco rattle virus (TRV)-mediated virus-induced gene silencing (VIGS). Chromatin immunoprecipitation (ChIP) assays further demonstrate that host proteins required for geminiviral replication — including the DNA polymerase δ subunits POLD1 and POLD3, RFC1, PCNA, topoisomerase I (TOPO1), and the single-stranded DNA-binding protein RPA70A—physically associate with viral DNA, supporting their role as components of the viral replisome. Importantly, Rep promotes the recruitment of RFC, POLD3, PCNA, TOPO1, and probably POLD1 to the viral genome. Our results indicate that geminiviruses co-opt components of the plant bidirectional replication fork to perform RCR; while lagging-strand-specific factors such as DNA ligase I (LIG1) and flap endonuclease 1 (FEN1) are dispensable for viral multiplication, suggesting leading strand-like replication, the DNA polymerases appear to be swapped, with DNA polymerase δ, normally associated with lagging-strand replication, mediating geminivirus DNA synthesis. Finally, we show that these host factors are required for replication across different geminivirus genera, as well as for a member of the CRESS *Nanoviridae* family. Together, these findings provide mechanistic insight into viral DNA replication and reveal vulnerable host dependencies that could be exploited for the rational design of antiviral resistance strategies.

## RESULTS

### The proxiome of the viral Rep in infected cells includes host-encoded DNA replication-related proteins

To gain insight into the composition of the geminiviral replisome, we performed TurboID-based PL, a technique that enables the detection of proteins in the vicinity of a protein of interest, including transient and weak interactors, using TYLCV Rep as bait. Because fusion tags can potentially alter the functional activity of the protein of interest, we first assessed the functionality of Rep fusion protein using a complementation assay. To this end, Rep fused to GFP or TurboID was transiently expressed in *Nicotiana benthamiana* leaves in the presence of a replication-deficient TYLCV Rep-null mutant. Our results showed that TYLCV Rep retained its ability to complement the Rep-null mutant only when fused to a C-terminal tag (GFP or TurboID), whereas this ability was largely abolished when the tag was fused to the N-terminus of Rep (Figure 1A, B). This observation may be attributed to the essential role of the N-terminal DNA binding/DNA nicking domain in viral DNA replication. Importantly, and consistent with its ability to functionally complement Rep-null mutant viruses, fusion to TurboID did not affect the nuclear localization of Rep (Figure 1C). In addition, Rep-TurboID retained biotinylation activity (Figure 1D, E).

**Figure 1.**
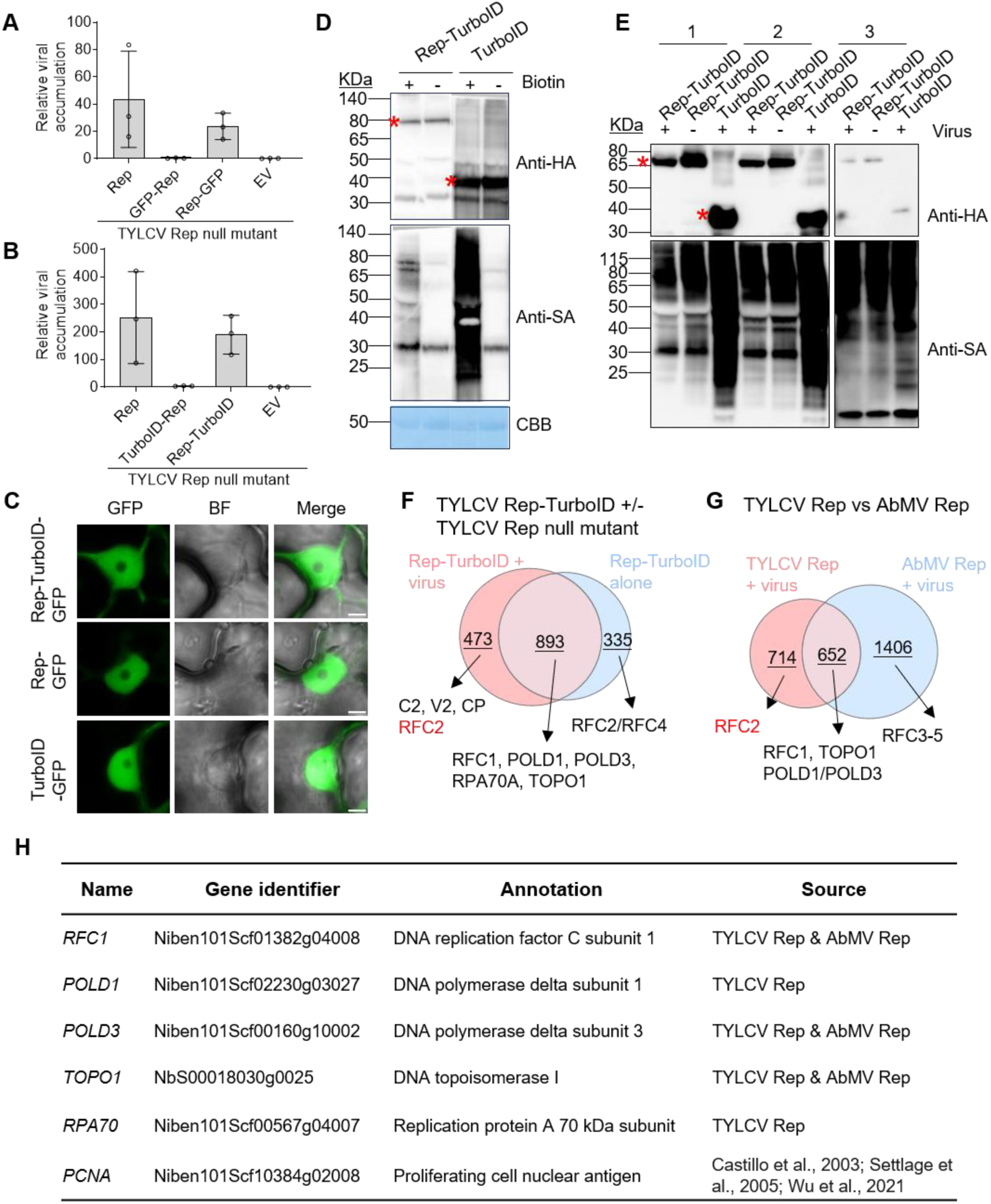
TYLCV Rep fused to a C-terminal TurboID moiety mediates viral genome replication and biotinylates DNA replication-related proteins. **A-B.** TYLCV Rep fused to a C-terminal tag (A: -GFP; B: -TurboID), retains its ability to mediate viral genome replication. *Agrobacterium* cells containing an infectious TYLCV clone harbouring a null mutation in Rep were co-infiltrated into *N. benthamiana* leaves with cells containing constructs to express Rep fusion proteins, untagged Rep (positive control), and empty vector (EV, negative control). Samples shown in (A) and (B) were harvested at 2 and 3 days post-infiltration, respectively. Viral DNA accumulation was measured by qPCR with 25S ribosomal DNA interspacer (ITS) as an internal reference. Experiments shown in (A) and (B) were repeated three times and twice, respectively, with similar results. **C.** Rep-TurboID localizes to the nucleoplasm. *Agrobacterium*-mediated transient expression of Rep-TurboID-GFP, Rep-GFP (positive control), and TurboID-GFP (negative control) in *N. benthamiana* plants. Confocal images were taken at 30 hours post-infiltration (hpi). BF, brightfield. Merge, overlay of fluorescence and BF images. Scale bar: 5 μm. This experiment was repeated twice with similar results. **D.** Immunoblot analysis of protein accumulation (anti-HA) and biotinylation activity (Streptavidin-HRP, anti-SA) in the presence and absence of exogenous biotin. At 42 hpi, *N. benthamiana* leaves were infiltrated with either 50 µM biotin (+Biotin) or DMSO (-Biotin) and harvested 6 hours later. CBB: Coomassie Brilliant Blue. This experiment was repeated twice with similar results. **E.** Immunoblot analysis of fusion protein level (anti-HA) and biotinylated proteins (anti-SA) in the samples from input lysates (1), after desalting but without affinity purification (2), and after affinity purification using streptavidin beads (3). Virus, TYLCV Rep null mutant. The position of asterisks indicates the expected bands of RT (∼78.0 kDa) and T (3xHA TurboID, ∼38.7 kDa), respectively. V, TYLCV Rep null mutant. This experiment was repeated three times with similar results. The positions of asterisks in panels D and E indicate the expected bands of Rep-TurboID (∼78.0 kDa) and TurboID (∼38.7 kDa), respectively. **F-G.** Venn diagram illustrates the overlap of Rep-labelled proteins in the presence and absence of TYLCV (F) and proteins labelled by TYLCV Rep and AbMV Rep in the presence of the corresponding virus (G). Arrows indicate known DNA replication-related proteins specifically labelled under different experimental conditions. Proteins shown in black meet the protein and peptide identification threshold of over 95%, while those labelled in red fall below this confidence threshold, as indicated. **H.** DNA replication-related proteins selected from the results in F and G for further study.

We next performed two parallel PL experiments: one with Rep-TurboID alone, and another one where Rep-TurboID complements the TYLCV Rep null mutant. Free TurboID served as negative control. Mass spectrometry (MS) data were filtered using the following criteria: (i) protein and peptide identification confidence ≥95%, and (ii) peptide counts in experimental samples at least two-fold higher than in control samples. The filtered datasets were visualized using Venn diagrams (Figure 1F; Figure S2A); the complete lists of identified factors are provided in Supplementary Tables 1 and 2. Thirteen proteins were consistently identified across all three biological replicates in TYLCV Rep samples in the context of infection (Figure S2A, Supplementary Table 1). Notably, four of these proteins are associated with RNA splicing (see Supplementary Table 1), consistent with recent studies demonstrating that geminiviruses manipulate the host RNA splicing machinery to facilitate infection (Pott *et al*., 2025, Preprint, 2026, Preprint). To obtain a comprehensive overview of potential Rep-labeled proteins, data from biological replicates were combined for subsequent analyses.

Several known host replication-associated proteins were identified among the putative TYLCV Rep-labeled host proteins under both experimental conditions, including RFC1, the catalytic (POLD1) and a regulatory (POLD3) subunits of Pol δ, TOPO1, and RPA70A. In the presence of the virus, three viral proteins, C2, V2, and CP, were detected as proximal to Rep, consistent with previous results (Wang *et al*., 2022).

To determine whether the identified replication-related factors play a conserved role among geminiviruses, we performed similar experiments with the Rep from AbMV, a bipartite begomovirus (Figure S1, Figure S2B, and Supplementary Tables 1-2). Surprisingly, in contrast to TYLCV Rep, AbMV Rep fused to TurboID at either the N- or C-terminus was able to complement the AbMV Rep-null mutant, although viral accumulation was reduced compared to the wild type (Figure S1A). This difference might be attributed to the longer N-terminal region of AbMV Rep, which could confer increased structural flexibility and allow proper folding upon N-terminal tagging (Figure S1B).

Similar results to those previously obtained with TYLCV Rep were observed with AbMV Rep, where RFC1-5, POLD1, POLD3, and TOPO1 were detected as proximal proteins. Cross-comparison of TYLCV and AbMV Rep proxiomes in the presence of their respective viruses revealed a shared core set of host factors, including multiple RFC subunits, Pol δ components, and TOPO1 (Figure 1G, Supplementary Tables 1-2).

Replication-associated proteins labelled by one or both Rep proteins were selected for further investigation (Figure S2 C-D). PCNA, the DNA clamp that serves as the processivity factor for DNA polymerase δ, has been shown to interact with geminivirus Rep (Castillo et al., 2003; Bagewadi et al., 2004). Because DNA polymerase δ is required for TYLCV replication (Wu et al., 2021) and two of its subunits as well as two subunits of the clamp loader RFC were identified as labelled by Rep, we decided to include PCNA in this selection.

All host replication-associated proteins selected for further characterization are listed in Figure 1H. Consistent with our proteomic results, all selected proteins localize to the nucleus, with TOPO1 and RFC1 showing enhanced nucleolar enrichment. RPA70A forms nuclear speckles, of which the number increases noticeably in the presence of either Rep or TYLCV (Figure S3). The association between these proteins and TYLCV Rep was confirmed by co-immunoprecipitation experiments (Figure S4).

Collectively, these results demonstrate that the Rep proximal proteome in two geminiviruses includes host DNA replication-related proteins, and that Rep physically associates with those factors.

### Rep proximal proteins are required for virus replication and components of the viral replisome

To assess the functional contributions of the selected DNA replication-related proteins to geminiviral genome replication, we employed TRV-mediated VIGS to knock-down the corresponding genes in *N. benthamiana* (Figure S5-S9). Importantly, *Agrobacterium tumefaciens*-mediated transient expression was not compromised in silenced plants (Figure S10).

Next, we evaluated the effect of silencing the selected host genes on Rep-mediated DNA replication by using either transgenic 2IR-GFP replication reporter *N. benthamiana* plants (Maio *et al*., 2019), in which GFP is overexpressed as a result of Rep DNA replication activity, or viral infection assays.

Silencing phenotypes were assessed at 14 days post-TRV inoculation, with the exception of *TOPO1*-silenced plants, which were observed and sampled at 7 days post-TRV inoculation, as silencing of this gene led to premature plant death (Figure 2A). Transient expression of TYLCV Rep in control 2IR-GFP transgenic *N. benthamiana* plants, inoculated with TRV2-*GUS*, led to a strong GFP signal, as previously described. However, the GFP signal was markedly reduced in *RFC1*-, *POLD1*-, *POLD3*-, *PCNA*-, and *TOPO1*-silenced 2IR-GFP plants (Figure 2B, C). In line with this observation, the accumulation of the IR-GFP replicon, resulting from Rep-dependent DNA replication on the reporter transgene, was markedly reduced in silenced 2IR-GFP plants (Figure 2D), indicating that the ability of Rep to mediate virus-like replication is impaired upon silencing of these genes. We next assessed the impact of gene silencing on local infections, where viral accumulation serves as a proxy for viral DNA replication (Wu *et al*., 2021). Viral accumulation was drastically reduced in all silenced plants compared to control plants (Figure S11; Figure 2E). The effect of silencing those replication-associated genes on inhibiting AbMV viral accumulation was similarly observed in AbMV locally infected plants (Figure S12); *TOPO1* was excluded from these analyses due to the severe phenotypic effects observed upon its silencing in *N. benthamiana* plants. Taken together, these results demonstrate that the selected replication-related proteins labelled by Rep play a critical role in the replication of two distinct begomoviruses.

**Figure 2.**
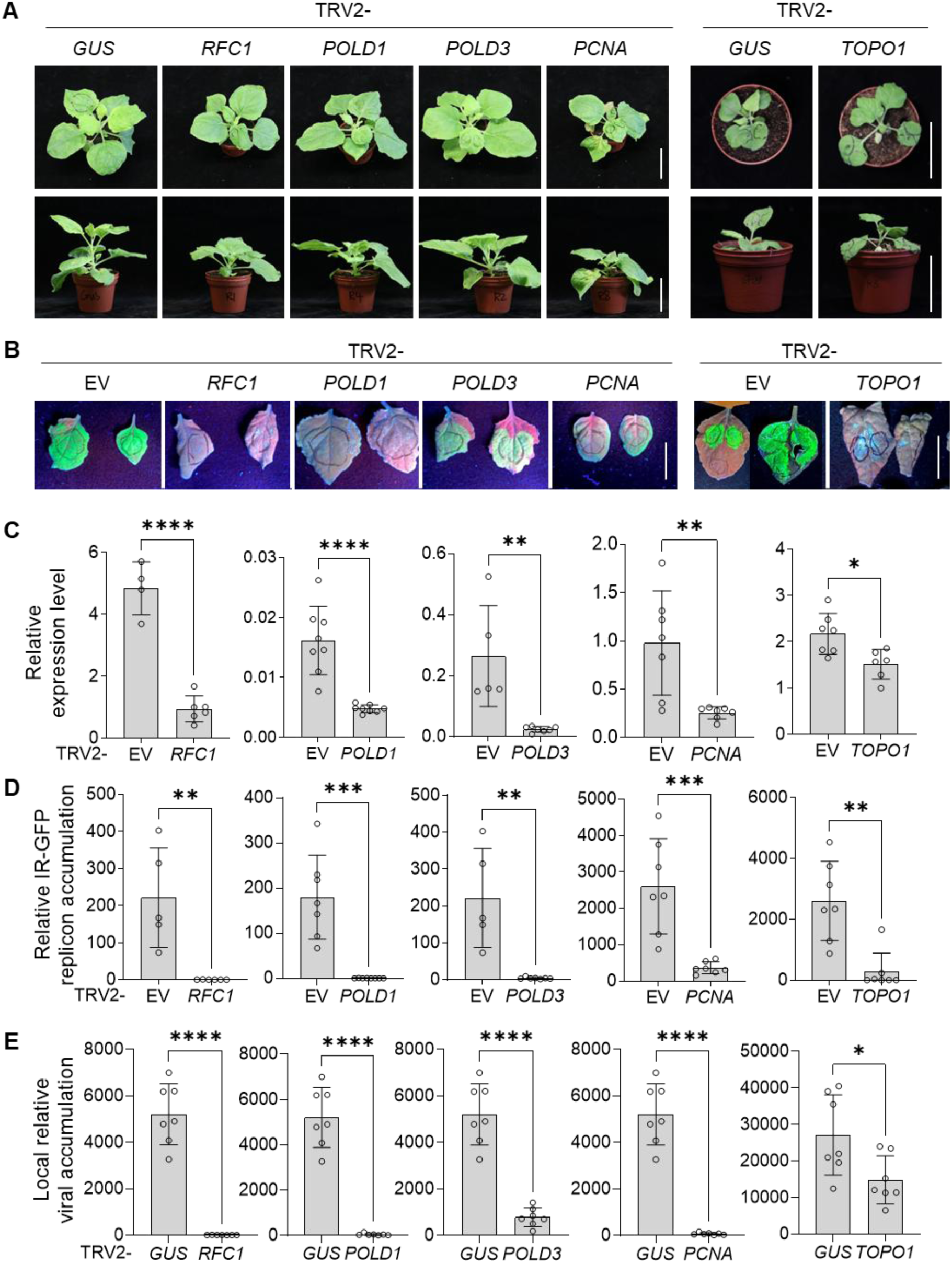
Selected replication-associated proteins are required for geminiviral replication. **A.** Developmental phenotypes of *RFC1*-, *POLD1*-, *POLD3*-, and *PCNA*-and *TOPO1*-silenced *N. benthamiana* plants. Plants infiltrated with TRV2-GUS were used as negative control. Scale bar: 5 cm. This experiment was repeated more than three times with similar results. **B-D.** GFP signal observation (B) and silencing efficiency (C) in *RFC1*-, *POLD1*-, *POLD3*-, *PCNA*-, and *TOPO1*-silenced 2IR-GFP transgenic *N. benthamiana* plants following agroinfiltration with Rep-RFP, measured by qRT-PCR using *NbActin* as an internal reference. (D) IR-GFP replicon accumulation measured by qPCR with 25S ribosomal DNA interspacer (ITS) as an internal reference. This experiment was repeated twice with similar results. **E.** Viral accumulation in *RFC1*-, *POLD1*-, *POLD3*-, *PCNA*-, and *TOPO1*-silenced wild-type (WT) *N. benthamiana* plants upon TYLCV local infection, measured by qPCR with 25S ribosomal DNA interspacer (ITS) as an internal reference. Samples expressing Rep-RFP (2IR-GFP transgenic plants) or inoculated with the TYLCV infectious clone (WT *N. benthamiana* plants) were harvested at 2 and 3 days post-infiltration, respectively. These experiments were repeated three times with similar results. Each dot represents an independent plant. Error bars indicate standard deviation. TRV2-EV or TRV2-GUS plants served as the control. Asterisks indicate statistically significant differences based on Student’s t-test (****, P<0.0001; ***, P<0.001; **, P<0.01; *, P<0.05; not significant (ns), P>0.05).

We then evaluated the impact of silencing these genes on TYLCV systemic infections. Consistent with milder symptoms observable in silenced plants, viral accumulation was significantly reduced (Figure 3A-C). Collectively, these findings indicate that the selected replication-related proteins are required for geminiviral infection, most likely due to their essential contribution to viral replication.

**Figure 3.**
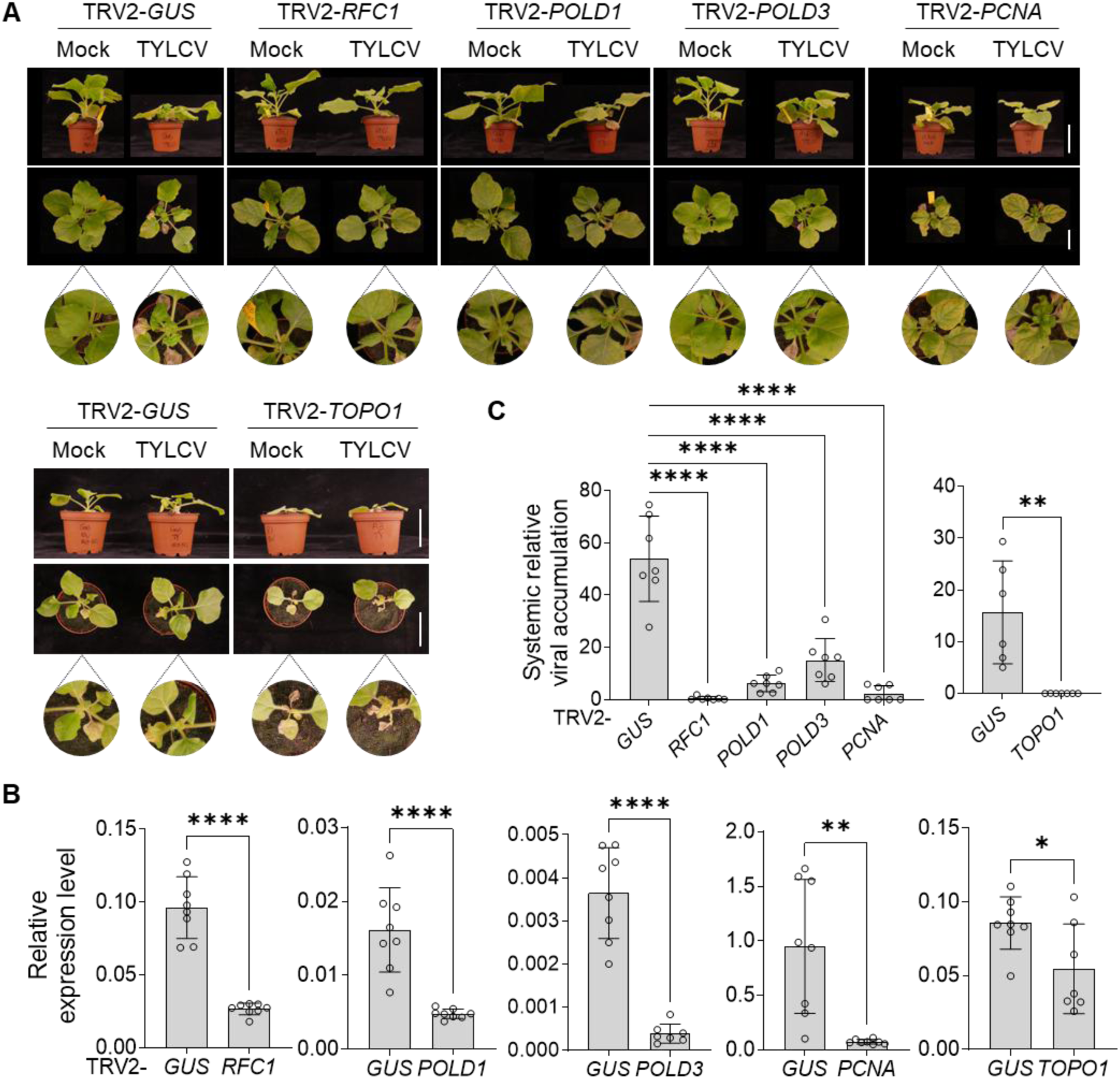
Selected replication-association factors are required for geminiviral infection. **A.** Symptom observation (A, scale bar: 5 cm) in *RFC1-*, *POLD1-*, *POLD3-*, *PCNA-*, and *TOPO1-*silenced *N. benthamiana* plants upon TYLCV systemic infection. **B.** Silencing efficiency measured by qRT-PCR using *NbActin* as an internal reference. **C.** Viral accumulation measured by qPCR with 25S ribosomal DNA interspacer (ITS) as an internal reference. *Agrobacterium* cells containing the TYLCV infectious clone, TRV1, and TRV2-RFC1, TRV2-POLD3, TRV2-POLD1, TRV2-PCNA, or TRV2-TOPO1 were mixed at a 1:1:1 ratio and inoculated into 2-week-old *N. benthamiana* seedlings. Samples were harvested at 14 days post-inoculation, except for *TOPO1-*silenced samples, which were harvested at 9 dpi. This experiment was repeated three times with similar results. Each dot represents an independent plant. Error bars indicate standard deviation. TRV2-GUS plants served as the control. Asterisks in panel B and on the right side of panel C indicate statistically significant differences based on Student’s t-test, whereas asterisks on the left side of panel C indicate statistically significant differences based on one-way ANOVA followed by Dunnett’s multiple comparisons test (****, P<0.0001; ***, P<0.001; **, P<0.01; *, P<0.05; not significant (ns), P>0.05).

We next sought to determine whether the selected proteins are components of the viral replisome. To achieve this, we examined their capacity to bind the viral genome using chromatin immunoprecipitation (ChIP) followed by quantitative PCR (ChIP-qPCR). To minimize protein competition issues, silencing-resistant (SR) versions of the target genes, harbouring silent mutations in the fragment used for VIGS that render them non-silenceable, were used to express FLAG-tagged proteins in silenced plants (Figure 4A, B). A construct to express Rep and either a construct to express the SR version of the silenced gene or an empty vector (EV) control were co-transformed into silenced 2IR-GFP transgenic *N. benthamiana* leaves at 14 days post-TRV inoculation. GFP signal was observed under UV light two days later. As shown in Figure 4B, the leaf patches co-infiltrated with Rep and the SR version exhibited a clearly stronger GFP signal compared to control patches. These results indicate successful complementation by the fusion proteins encoded by the SR gene versions in the silenced *N. benthamiana* background, demonstrating that the negative effect observed on viral DNA replication (Figure 2) is indeed specifically due to the knockdown of the respective target genes. We then performed ChIP-qPCR to assess the binding of these FLAG-tagged replication-associated proteins to the IR-GFP replicon in 2IR-GFP transgenic plants. The positions of the amplified sequences are shown in Figure 4C. Our results indicate that all five replication-associated proteins, RFC1, POLD1, POLD3, PCNA, and TOPO1, can physically bind to the IR-GFP replicon. Despite our inability to successfully silence *RPA70A*, ChIP-qPCR results indicate that RPA70 can also be associated with the replicon (Figure 4C). Binding of these factors to different regions of the viral genome can also be detected in local infection assays (Figure 4D).

**Figure 4.**
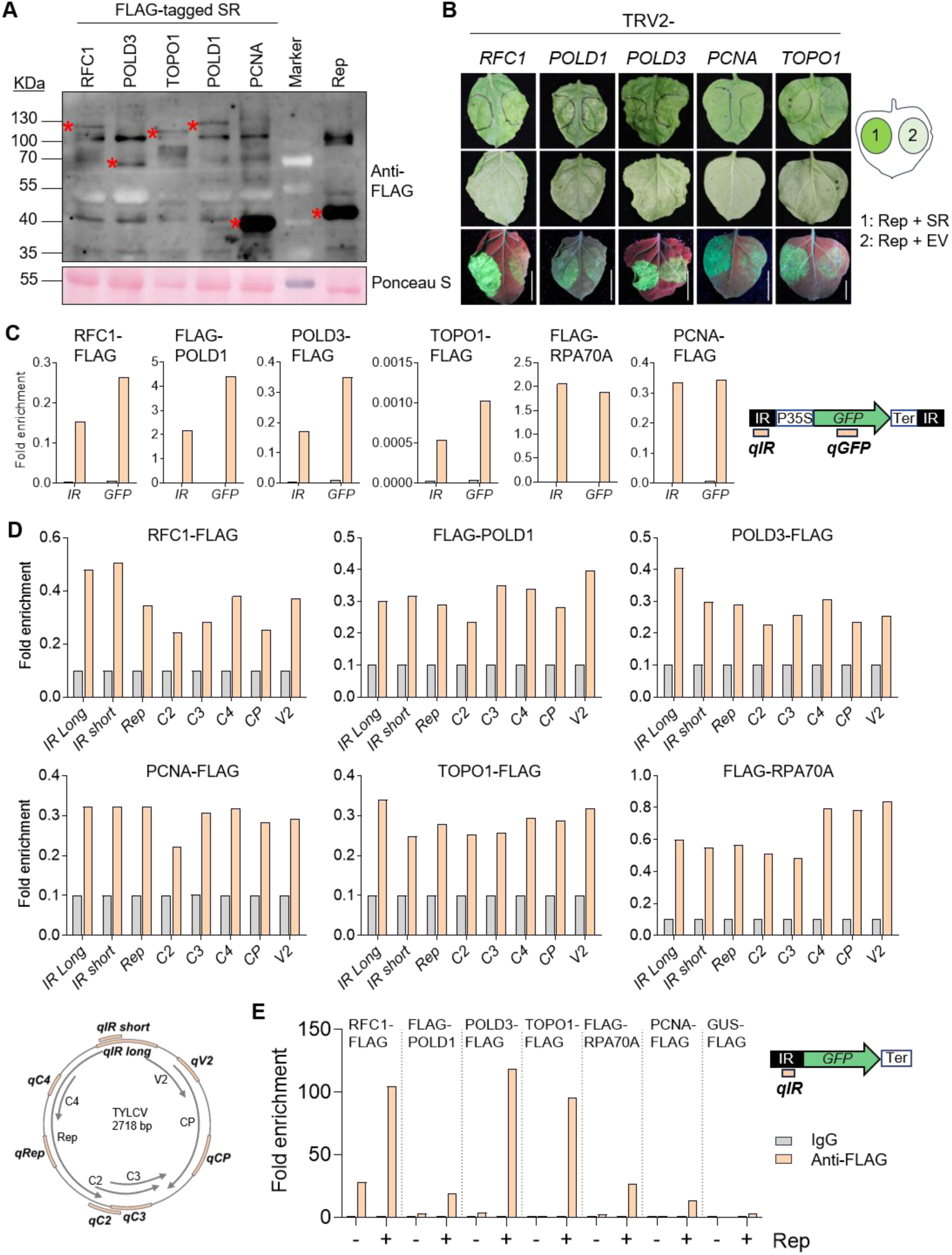
Selected replication-associated factors are likely components of the geminiviral replisome. **A.** Western blot analysis of the accumulation of silencing-resistant (SR)-encoded replication-associated proteins. Rep-FLAG was used as the positive control. Ponceau S, ponceau staining. The predicted protein sizes are as follows: RFC1-FLAG-SR, ∼106 kDa; POLD3-FLAG-SR, ∼57 kDa; TOPO1-FLAG-SR, ∼97 kDa; FLAG-POLD1-SR, ∼120 kDa; PCNA-FLAG-SR, ∼30 kDa; Rep-FLAG, ∼40 kDa. Red asterisks indicate the expected bands. **B.** Functional analysis of fusion proteins encoded by the SR gene versions in *RFC1-*, *POLD1-*, *POLD3-*, *PCNA-*, and *TOPO1*-silenced 2IR-GFP *N. benthamiana* plants. Constructs to express FLAG-tagged proteins or the corresponding empty vector (EV) and Rep were transiently transformed in gene-silenced 2IR-GFP transgenic *N. benthamiana* plants. GFP signal was observed at 2 days post-infiltration under a UV-lamp. Scale bar: 2 cm. **C-D.** RFC1, POLD1, POLD3, PCNA, TOPO1, and RPA70A bind to the IR-GFP replicon in the 2IR-GFP reporter system (C) and to the viral genome upon viral infection (D). Transient expression of FLAG-tagged proteins and Rep or the TYLCV infectious clone in gene-silenced 2IR-GFP transgenic (C) or wild-type *N. benthamiana* plants (D), respectively. All constructs are SR with the exception of RPA70A, which was overexpressed in the non-silenced background. All experiments were repeated at least twice with similar results. **E.** RFC1, POLD3, TOPO1, and PCNA bind the geminiviral IR region in a Rep-dependent manner. Transient transformation of constructs to express RFC1-FLAG/FLAG-POLD1/POLD3-FLAG/TOPO1-FLAG/FLAG-RPA70A/PCNA-FLAG or GUS-FLAG (negative control) with constructs to express Rep-RFP (+ Rep) or RFP (-Rep) and IR-GFP in 4-week-old WT *N. benthamiana* plants. All experiments were repeated at least three times with similar results. Schematic representations of the IR-GFP replicon (C), the viral genome (D), and the IR region (E) are shown at the bottom of panel D and to the right of panels C and E, respectively. The positions of the ChIP-qPCR primer pairs are indicated in light orange. Grey and light orange boxes represent IgG (negative control) and anti-FLAG, respectively.

Rep is believed to recruit the host DNA replication machinery to the viral DNA. To test whether Rep is required for binding of these host proteins to the viral origin of replication, we evaluated their capacity to bind the intergenic region (IR) of the viral genome (containing the origin of replication) in the presence or absence of this virus-encoded protein. As shown in Figure 4E and Figure S13, RFC1, POLD3, TOPO1, RPA70A, and PCNA, as well as, to a lesser extent, POLD1, displayed a clear enrichment in IR binding in the presence of Rep in at least two out of three independent biological replicates, indicating that Rep recruits these factors to the viral DNA, either directly or indirectly.

In summary, our results demonstrate that Rep-labelled proteins required for viral replication and infection are likely components of the viral replisome, and their association with the viral genome is dependent on the presence of Rep.

### The geminiviral genome likely follows leading strand-like replication mediated by the lagging strand-associated DNA polymerase δ

Unlike eukaryotic chromosomal replication, which proceeds through bidirectional replication forks, geminiviral RCR is unidirectional, suggesting that these viruses employ a distinct genome replication strategy from that of their hosts. This fundamental difference raises the question of whether geminiviruses rely on leading-strand-like synthesis, lagging-strand-like synthesis, or a combination of both. Because putative geminiviral replisome components identified above (PCNA, RFC, RPA, and TOPO1) participate in the replication of both DNA strands, we investigated whether the lagging-strand replication machinery contributes to geminiviral replication by examining the involvement of the specific components FEN1, which removes 5′ flap intermediates generated during lagging-strand synthesis, and LIG1, which seals the resulting DNA nicks to complete Okazaki fragment maturation. For this purpose, *FEN1* and *LIG1* were silenced by VIGS (Figures S14-S17; Figure 5) in 2IR-GFP or wild-type *N. benthamiana* plants, and the effect of their silencing on Rep-mediated replication or geminiviral infection was evaluated. Despite efficient silencing of FEN1 (Figure S18), no impact on GFP replicon, or virus accumulation, either in local or in systemic infections, could be detected (Figure 5; Figure S19).

**Figure 5.**
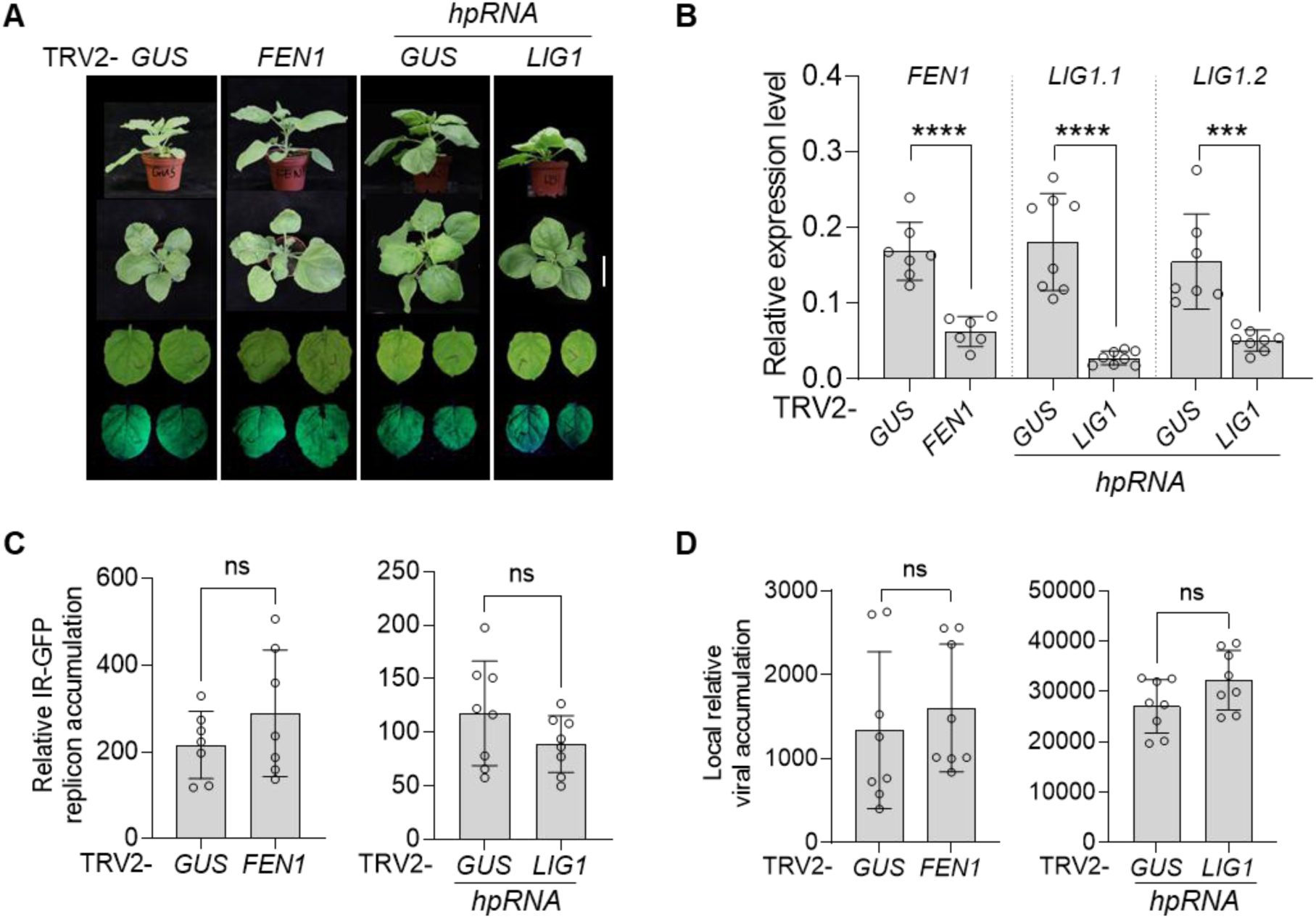
Lagging strand-specific DNA replication-related proteins are dispensable for Rep-dependent DNA replication. **A.** Developmental phenotypes (top panel) and GFP signal (bottom panel) of *FEN1*- and *LIG1*-silenced 2IR-GFP transgenic *N. benthamiana* plants following transient expression of Rep-RFP. Scale bar: 5 cm. **B-C.** Silencing efficiency (B) of the plants in (A), measured by qRT-PCR using *NbActin* as an internal reference, and IR-GFP replicon accumulation (C), measured by qPCR with 25S ribosomal DNA interspacer (ITS) as an internal reference. This experiment was repeated twice with similar results. **D.** Viral accumulation in *FEN1*- and *LIG1*-silenced wild-type (WT) *N. benthamiana* plants upon TYLCV local infection, measured by qPCR with 25S ribosomal DNA interspacer (ITS) as an internal reference. Plants infiltrated with TRV2-GUS were used as negative control. For LIG1, the hairpinRNA-*NbLIG1.2* construct was transiently co-transformed with either that to express Rep-RFP in *NbLIG1.1*-silenced 2IR-GFP transgenic plants or with the TYLCV infectious clone in *NbLIG1.1*-silenced wild-type plants at 14 dpi. Samples expressing Rep-RFP or inoculated with the TYLCV infectious clone were harvested at 2 and 3 days post-infiltration, respectively. Each dot represents an independent plant. Error bars indicate standard deviation. TRV2-GUS plants served as the control. Experiments shown in panels A–C were repeated three times for FEN1 and twice for LIG, whereas the experiment shown in panel D was repeated twice for FEN1 and three times for LIG1, with similar results. Asterisks indicate statistically significant differences based on Student’s t-test (****, P<0.0001; ***, P<0.001; **, P<0.01; *, P<0.05; not significant (ns), P>0.05).

The VIGS construct used for *LIG1* is predicted to efficiently knock down only one of the two *N. benthamiana LIG1* isoforms, *NbLIG1.1* (Figure S14). Therefore, a hairpin RNA-mediated silencing approach was employed to simultaneously target *NbLIG1.2*. To this end, the hpRNA-*NbLIG1.2* construct was transiently co-expressed with either Rep or the TYLCV infectious clone in *NbLIG1.1*-silenced plants at 14 dpi, and samples were collected 2–3 days later for further analyses. As shown in Figure 5 and Figure S18, silencing of both *LIG1* homologues does not produce any effect on replicon or viral accumulation, suggesting that, as FEN1, LIG1 is not required for Rep-dependent DNA replication.

The dispensability of FEN1 and LIG1 for Rep-dependent DNA replication suggests that geminiviral RCR likely proceeds via a continuous, leading strand-like mechanism, rather than classical lagging-strand synthesis requiring Okazaki fragment processing. Intriguingly, it is DNA polymerase δ, which mediates mostly lagging-strand replication in the eukaryotic bidirectional replication fork, the one involved in this process (Wu *et al*., 2021). Collectively, these findings provide experimental support for a leading-strand-driven RCR model for geminiviruses, involving recruitment of host leading-strand machinery with a swap in DNA polymerases. Of note, a precedent for non-canonical strand/polymerase assignment exists in the replication of the polyomavirus simian virus 40 (SV40), which is thought to rely predominantly on host DNA polymerase δ for synthesis of both the leading and lagging strands (Tsurimoto and Stillman, 1991a,b).

### A subset of host DNA replication factors is commonly required for replication of CRESS DNA viruses

CRESS DNA viruses ubiquitously employ RCR to multiply their genomes. To gain deeper insight into the conservation of host factors involved in CRESS DNA viral replication, we conducted a comparative analysis across different viral species. First, we assessed the effect of silencing selected DNA replication-associated genes, found to be required for TYLCV and AbMV replication (*RFC1*, *POLD1*, *POLD3*, and *PCNA*), on the replication of chickpea chlorotic dwarf virus (CpCDV), a member of the *Mastrevirus* genus within the family *Geminiviridae* (Figure S20).

Similarly, in the case of CpCDV, as previously observed for the begomoviruses TYLCV and AbMV, local viral accumulation, reflecting mostly replication, was almost completely suppressed in silenced plants (Figure 6A, B). As expected, systemic infections were also affected, with a siginificant reduction in viral accumulation and symptom development (Figure 6C and D; Figure S21A). Taken together, these results demonstrate the essential roles of replication-associated proteins labelled by TYLCV and AbMV Rep in PL assays in mastrevirus replication. Considering the phylogenetic divergence between CpCDV and TYLCV (Figure S20A), these results support the notion that the function of these replication-related proteins in geminiviral replication is likely conserved across different geminiviral species.

**Figure 6.**
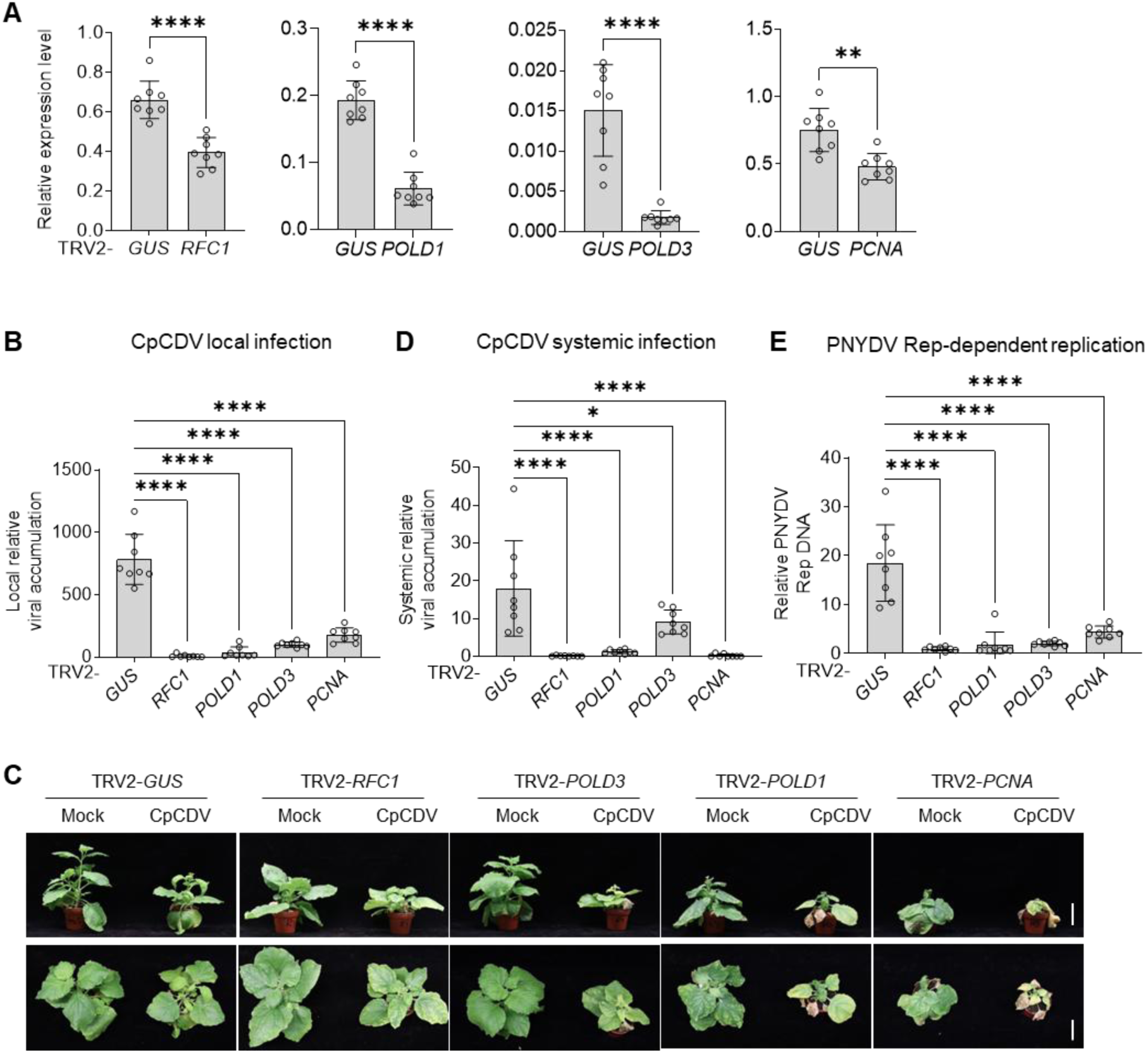
Components of the eukaryotic DNA replication fork are required for replication of CRESS viruses. A-B. Silencing efficiency (A) in *RFC1*-, *POLD1*-, *POLD3*-, and *PCNA*-silenced *N. benthamiana* plants upon CpCDV local infection, measured by qRT-PCR using *NbActin* as an internal reference, and viral accumulation (B), measured by qPCR with 25S ribosomal DNA interspacer (ITS) as an internal reference. Samples inoculated with the CpCDV infectious clone were harvested at 3 days post-infiltration. **C-D.** Symptoms (C, scale bar: 5cm) in *RFC1*-, *POLD1*-, *POLD3*-, and *PCNA*-silenced *N. benthamiana* plants upon CpCDV systemic infection. Viral accumulation (D), measured by qPCR with 25S ribosomal DNA interspacer (ITS) as an internal reference. *Agrobacterium* cells containing the CpCDV infectious clone, TRV1, and TRV2-RFC1, TRV2-POLD3, TRV2-POLD1, or TRV2-PCNA were mixed at a 1:1:1 ratio and inoculated into 3-week-old *N. benthamiana* seedlings. Samples were harvested at 14 days post-inoculation. **E.** PNYDV DNA-R quantification in *RFC1*-, *POLD1*-, *POLD3*-, and *PCNA*-silenced *N. benthamiana*, measured by qPCR with 25S ribosomal DNA interspacer (ITS) as an internal reference. Samples infiltrated with the PNYDV DNA-R component were harvested at 2 days post-infiltration. Each dot represents an independent plant. Error bars indicate standard deviation. TRV2-GUS plants served as the control. Asterisks in panel A indicate statistically significant differences based on Student’s t-test, whereas asterisks in panels B, D and E indicate statistically significant differences based on one-way ANOVA followed by Dunnett’s multiple comparisons test (****, P<0.0001; ***, P<0.001; **, P<0.01; *, P<0.05; not significant (ns), P>0.05). All experiments were repeated at least twice with similar results.

*Nanoviridae* is a second family of plant-infecting viruses belonging to the CRESS DNA phylum (Figure S20B). Unlike most geminiviruses, nanoviruses have multipartite genomes, consisting of 6-8 circular ssDNA molecules of approximately 1 kb in size (Lal *et al*., 2020). Similar to geminiviral Rep, the Rep protein encoded by nanoviruses is the only viral protein essential for viral DNA replication. To further investigate whether host factors labelled by geminiviral Rep are also required for replication of other CRESS DNA viruses, we examined their roles in DNA replication mediated by the Rep protein of the nanovirus pea necrotic yellow dwarf virus (PNYDV). Because the Rep-encoding DNA-R component from some nanoviruses has been reported to replicate autonomously in *N. benthamiana* (Timchenko *et al*., 1999), we hypothesized that the PNYDV DNA-R component would behave similarly. Therefore, *A. tumefaciens*-mediated delivery of the circular PNYDV DNA-R replicon into silenced *N. benthamiana* plants allowed us to assess the role of geminiviral Rep-labelled proteins in PNYDV

Rep-dependent replication. As shown in Figure 6E and Figure S21B, PNYDV Rep DNA levels were detectable in wild-type *N. benthamiana* plants, indicating replication of DNA-R, but dramatically decreased in silenced plants. Collectively, these findings suggest that RFC1, POLD1, POLD3, and PCNA are essential for replication of nanoviruses.

Overall, our results are in line with a model in which CRESS DNA viruses selectively co-opt core host replication factors common to both leading- and lagging-strand synthesis, while likely bypassing lagging strand-specific processing components and swapping the DNA polymerases. This implies a conserved replication strategy among CRESS DNA viruses, wherein selected host factors involved in eukaryotic DNA replication are recruited by CRESS DNA virus Rep proteins and repurposed to perform RCR (Figure 7).

**Figure 7.**
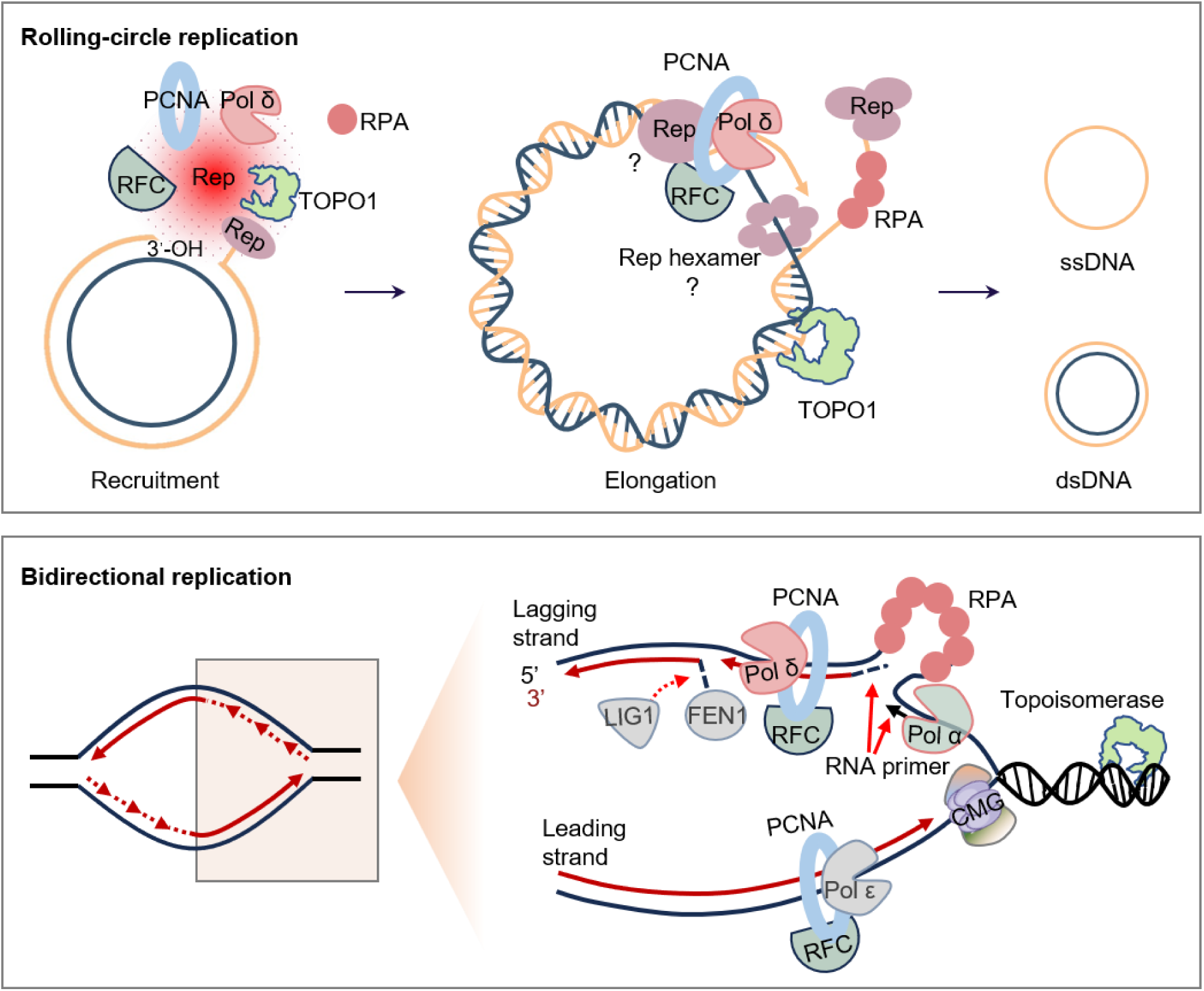
Proposed molecular events underlying geminiviral rolling-circle replication (RCR). The double-stranded DNA (dsDNA) replicative intermediate, generated from the viral single-stranded DNA (ssDNA) genome, serves as the template for RCR. The viral Rep protein specifically recognizes and nicks the conserved nonanucleotide sequence within the origin of replication on the virion strand, initiating DNA synthesis. Rep likely unwinds the viral DNA duplex as a hexamer, whereas lower-order Rep oligomers may selectively recruit components of the host bidirectional DNA replication machinery, including TOPO1, RFC, PCNA, DNA polymerase δ, and RPA, but not the canonical lagging-strand maturation factors FEN1 and LIG1. These recruited factors together form the viral replisome and mediate DNA strand elongation during RCR. As a result, the initial virion strand is released from the dsDNA replicative form. For simplicity, the initial conversion of the viral ssDNA genome into the dsDNA replicative form is not depicted. Orange and dark blue indicate the virion and complementary DNA strands, respectively.

## DISCUSSION

Viral genome replication is central to infection and therefore represents an attractive target for interventions aimed at achieving durable antiviral resistance. Geminiviruses rely almost entirely on the host DNA replication machinery for this process, with the virus-encoded Rep protein constituting the only viral factor essential for replication of their DNA genomes within the nucleus of infected cells. Despite the presumed parallels with eukaryotic DNA replication and the conceptual and mechanistic similarities to extensively studied animal DNA tumor viruses, our understanding of the molecular events governing geminiviral replication remains limited.

A major obstacle to the study of geminiviral replication is that many host factors involved in DNA replication are essential genes, precluding the generation of conventional loss-of-function mutant plant lines. In addition, experimental approaches to investigate viral replication *in vivo* are scarce. Here, we overcame these challenges by utilizing VIGS for the functional interrogation of candidate host factors derived from TurboID-based PL of Rep in complementation assays performed *in planta*. PL by replication-competent, TurboID-fused geminiviral Rep proteins (TYLCV and AbMV) during infection yielded a first comprehensive catalog of Rep-proximal proteins, including numerous components of the host DNA replication machinery.

From the list of putative Rep-proximal proteins, we selected several factors as proof of concept. These included host proteins with established roles in eukaryotic genome replication, some of which had previously been reported to associate with geminiviral Rep proteins through protein–protein interactions (Luque *et al*., 2002b; Castillo *et al*., 2003; Bagewadi *et al*., 2004) (Figure 1). Our results demonstrated that all selected factors i) are required for Rep-dependent DNA replication, ii) are essential for geminiviral replication and infection, and iii) bind the viral genome, supporting their role as likely components of the geminiviral replisome. Notably, the vast majority of host proteins identified within the Rep proximal proteome remain uncharacterized in the context of geminivirus infection. Given the results obtained thus far, we anticipate that this dataset will provide a valuable resource for advancing our understanding of geminivirus biology and of the multiple functions performed by Rep during infection.

The investigated replication-associated factors (RFC, PCNA, TOPO1, RPA) are involved in leading- and lagging-strand DNA replication. To discern whether both modes of replication are required for Rep-dependent geminivirus replication, we tested two proteins exclusively involved in the replication of the lagging strand, namely FEN1 and LIG1. The apparent dispensability of these two factors is consistent with the notion that geminiviruses utilize a predominantly leading strand-like replication mode. These results led us to suggest that geminiviruses, and likely other CRESS DNA viruses, selectively recruit host factors, including RFC, PCNA, RPA, and TOPO1, to their genomes, either directly or indirectly, with leading strand-like replication ensuing. Despite the accepted involvement of DNA polymerase δ in lagging-strand synthesis, this is the DNA polymerase involved in geminiviral replication, and the only main replicative polymerase labelled by Rep (Wu *et al*., 2021). Of note, DNA polymerase δ can mediate leading-strand replication during double-strand break repair in yeast, a situation in which it becomes error-prone (Hicks *et al*., 2010). In combination with PCNA, DNA polymerase δ is highly processive in the absence of downstream Okazaki fragments and, therefore, could replicate a complete geminivirus genome in one replication step. This inherent high processivity may be taken advantage of by geminiviruses to efficiently perform RCR (Langston and O’Donnell, 2008).

Based on our results, we propose the following sequence of molecular events underlying geminiviral Rep-dependent RCR, which utilizes a dsDNA intermediate converted from the single-stranded viral genome as template (Figure 7). At the onset of RCR, Rep selectively co-opts core host replication factors to the viral genome. Following nicking at the origin of replication by Rep, a 5’ end and a 3’ hydroxyl group are generated. The 5′ end is covalently bound by Rep (Laufs *et al*., 1995), while the 3′ end serves as a primer for extension by Pol δ, assisted by the sliding clamp PCNA, which is loaded onto the viral DNA by the clamp loader RFC. As replication proceeds, TOPO1, also directly or indirectly recruited by Rep, functions on alleviating torsional stress ahead of the replication fork. The exposed ssDNA generated during replication is coated by RPA, preventing degradation and secondary structure formation. At the termination of RCR, Rep catalyzes the re-ligation of the original nick introduced at the initiation step, resulting in the production of a copy of the initial ssDNA molecule, which is released from the circular dsDNA genome; this new ssDNA can re-enter the replication cycle or be transported (intra- and inter-cellularly or long-distance) or encapsidated. Given the requirement of some of the investigated factors for the replication of diverse geminiviruses as well as a nanovirus, this model might generally apply to CRESS DNA viruses across hosts.

Interestingly, DNA polymerase δ seems to be preferentially co-opted by independently evolved DNA viruses replicating in the nucleus of infected cells. The dependency of viral replication on this polymerase has been shown not only for geminiviruses (Wu *et al*., 2021) and nanoviruses (this work), but also for the animal-infecting CRESS DNA virus porcine circovirus (PCV), a member of the family *Circoviridae* (Gassmann *et al*., 1988). Beyond CRESS DNA viruses, replication of parvoviruses (family *Parvoviridae*), which have linear ssDNA genomes, occurs via a rolling-hairpin replication mode, considered a variation of RCR, and also relies on DNA polymerase δ (Nash *et al*., 2008). SV40 (family *Polyomaviridae*) has a dsDNA genome that replicates bidirectionally, but utilizes DNA polymerase δ for replication of both strands (Tsurimoto and Stillman, 1991a,b). Importantly, both SV40 and CRESS viruses provide DNA helicase for viral replication. In eukaryotes, the MCM (Minichromosome Maintenance) helicase complex tracks along the leading strand in association with DNA polymerase ε, which depends on this specific enzyme for DNA synthesis. DNA polymerase δ, on the other hand, is not associated with a specific host DNA helicase, and interacts with SV40 T antigen to mediate leading-strand replication. Association of Rep, which has been shown to possess DNA helicase activity (Choudhury *et al*., 2006; Clérot and Bernardi, 2006; George *et al*., 2014; Ruhel *et al*., 2021), with this polymerase might also facilitate coordination of template unwinding and RCR. The ability of DNA polymerase δ to interact with diverse helicases may make it the default choice for most viruses encoding with their own helicase enzymes. Whether these viruses exploit the increased mutation rate exhibited by Pol δ during certain DNA replication processes to promote their evolutionary adaptability remains an enticing question for future investigation. Intriguingly, preferential utilization of DNA polymerase δ has also been associated with several non-canonical DNA synthesis processes, including transposon-associated repair replication and break-induced replication, suggesting that reassignment of strand/polymerase usage may represent a broader feature of recombination- or strand displacement-driven DNA replication mechanisms (Fuchs *et al*., 2021).

Understanding the mechanisms underlying geminiviral genome replication provides a valuable framework for the development of antiviral strategies aimed at impairing viral multiplication while minimizing the potential for viral adaptation, thereby increasing the likelihood of durable resistance. Recent studies demonstrate the feasibility of such approaches. For example, nonsynonymous single-nucleotide polymorphisms (SNPs) in POLD1 (catalytic subunit of DNA polymerase δ) and PRIM2 (primase large subunit of DNA polymerase α) have been associated with resistance to geminiviruses in tomato and melon, respectively (Siskos *et al*., 2023; Shen *et al*., 2025). POLD1 variant alleles have been proposed to confer resistance across a broad range of crop species (Gilbert *et al*., 2025), including cassava (Lim et al., 2022), although this is currently under debate (Cornet et al., 2025). In this context, targeted disruption of the functional interplay between geminiviral Rep and the host DNA replication machinery may represent an effective strategy to achieve broad and durable resistance against these viruses.

Collectively, our findings not only provide long-sought mechanistic insight into geminiviral replication, but also establish a foundation for the rational development of antiviral strategies against geminiviruses and potentially other DNA viruses that exploit host DNA replication pathways.

## METHODS

### Plant material and bacterial and viral strains

Wild-type (WT) and 2IR-GFP transgenic *N. benthamiana* plants were grown in a controlled growth room under long-day conditions (16 hours light/8 hours dark) at 22 °C and 35% humidity.

*Escherichia coli* strain TOP10 was used for general cloning and subcloning, whereas *E. coli* strain DB3.1 was used for amplification of Gateway-compatible empty vectors. *Agrobacterium tumefaciens* strain GV3101 harbouring the corresponding binary vectors was used for *in planta* transformation.

The viruses and strains were TYLCV Almería isolate (TYLCV-Alm, GenBank accession no. AJ489258); AbMV-A and AbMV-B (GenBank accession no. NC_001928 and NC_001929); CpCDV (GenBank accession no. Y11023.2); PNYDV-R (GenBank accession no. GU553134).

### Plasmid construction

Plasmids and primers used for cloning are summarized in Supplementary Tables 3 and 4, respectively. All constructs in this study were generated using one of two cloning methods: Gateway cloning (Nakagawa *et al*., 2007a,b) was used when cloning into plasmids of the pGWB series. Traditional restriction/ligation cloning was used when cloning into the pTRV2 vector. DNA fragments cloned into the pDONR™/Zeo entry vector (Thermo Scientific) were recombined into the corresponding destination vectors through a Gateway LR reaction (Thermo Scientific).

### *A. tumefaciens*-mediated transient transformation

*A. tumefaciens* cells carrying the desired constructs were cultured overnight in LB medium supplemented with the appropriate antibiotics at 28 °C. Cells were harvested by centrifugation at 4,000 × g for 10 min at room temperature and resuspended in infiltration buffer (10 mM MgCl₂, 10 mM MES, pH 5.6, and 150 μM acetosyringone). The bacterial suspension was adjusted to an OD600 of 0.1–0.6 and incubated in the dark for 2–4 h before infiltration. *A. tumefaciens* suspensions were infiltrated into the abaxial side of leaves using a needleless syringe. For co-infiltration experiments, *A. tumefaciens* cultures were mixed at a 1:1 ratio before infiltration. Following infiltration, plants were kept in the growth room for 24–48 h before further analyses.

### Protein extraction and co-immunoprecipitation (co-IP) assays

*A. tumefaciens* cells carrying the desired constructs were infiltrated into fully expanded leaves of four-week-old *N. benthamiana* plants and were sampled at two days post-infiltration. The samples were ground into a fine powder with liquid nitrogen. For proteins with exclusive nuclear localization, Honda buffer was used to extract the nuclei. Protein extraction and co-IP with GFP-Trap agarose beads (Chromotek, GTA-20) were performed as described by Wu *et al*. (2021).

The antibodies used for co-IP assays were anti-GFP (SICGEN, AB0020-500; dilution ratio: 1:5000), anti-RFP (Chromotek, 6G6; dilution ratio: 1:5000), anti-goat IgG (Sigma, A8919-2ML; dilution ratio: 1:15,000), and anti-mouse IgG (Sigma, A2554; dilution ratio: 1:15,000).

### Visualization of protein subcellular localization

Confocal microscopy was used to visualize the subcellular localization in transiently transformed *N. benthamiana* leaves. *A. tumefaciens* cells expressing GFP- or RFP-tagged proteins were mixed at a 1:1 ratio and infiltrated into *N. benthamiana* leaves as described above. The samples were collected at 30-48 h post-infiltration and imaged with a Zeiss LSM 880 upright confocal microscope. The preset settings for GFP (Ex: 488 nm, Em: 500–550 nm) and RFP (Em: 570–620 nm) were used for imaging.

### Virus-induced gene silencing (VIGS)

Tobacco rattle virus (TRV)-mediated virus-induced gene silencing (VIGS) assays were performed as described by Wu *et al*. (2021). In brief, *A. tumefaciens* cells containing pTRV1 (helper plasmid), pTRV2-GUS (negative control), or constructs cloned in pTRV2 (Supplementary Table 3) were mixed and infiltrated into the leaves of two-week-old 2IR-GFP or wild-type *N. benthamiana* seedlings using a 1 mL needleless syringe. Pants were then kept in the growth room for 14 days for further analyses, except for *TOPO1*-silenced plants, which were kept for 7 days to avoid plant death.

### Local and systemic infections

Local and systemic viral infections were performed as described by Wu *et al*. (2021). Briefly, for local infection, the *A. tumefaciens* cells containing the infectious clones were infiltrated into wild-type *N. benthamiana* plants and samples were harvested at three days post-inoculation. For systemic infection, *A. tumefaciens* cells containing geminivirus infection clones were infiltrated into the leaves of two-week-old *N. benthamiana* seedlings. Samples were harvested at 14 days post-inoculation.

### Quantitative real-time PCR (qPCR) and reverse transcription PCR (RT-qPCR)

DNA and RNA were extracted as described by Oñate-Sánchez and Vicente-Carbajosa (2008). To detect gene expression in *N. benthamiana* plant tissue, 5 μg of crude RNA were digested with DNase I (Thermo Scientific) to remove genomic DNA. cDNA synthesis was then performed using PrimeScript RT Master Mix (Takara) following the manufacturer’s instructions. *NbActin* was used as the reference gene (Maimbo *et al*., 2010). For DNA samples from local viral infections, *Dpn*I (Thermo Scientific) was used to digest plasmid DNA potentially derived from *A. tumefaciens* prior to qPCR. The 25S ribosomal DNA interspacer (*ITS*) was used as the reference sequence (Mason *et al*., 2008). PowerTrack™ SYBR Green Mastermix (Thermo Scientific) was used for the qPCR assay in a BioRad CFX384 real-time system. The procedure was performed as follows: initial denaturation at 95 °C for 3 minutes, followed by 40 cycles of 95 °C for 15 seconds and 60 °C for 30 seconds. At least two technical replicates were performed for each experiment. qPCR primers were designed using the online tool Primer 3 and are shown in Supplementary Table 4.

### Chromatin immunoprecipitation (ChIP) assay

ChIP assays were performed as described by Nie *et al*. (2019) with minor modifications. *A. tumefaciens* cells carrying the desired constructs to express FLAG-tagged fusion proteins (as shown in Supplementary Table 3) were co-infiltrated with cells carrying constructs to express Rep-RFP or harbouring the TYLCV infectious clone into 2IR-GFP transgenic or WT *N. benthamiana* plants. Samples were harvested at two days post-infiltration. Crosslinking was performed using 1% formaldehyde in 1 x PBS by vacuum infiltration. The crosslinking reaction was then quenched with 0.125 M glycine in 1 x PBS. The following steps were carried out as described by Nie *et al*. (2019).

For the IP step, 2 µg of Anti-FLAG (SICGEN) or IgG (Sigma) were used. DNA was purified using the MinElute PCR Purification Kit (QIAGEN, No. 28004). ChIP input and IP products were diluted to a 1:1000 and 1:10 ratio, respectively. 1 µL was taken for qPCR analysis. The primers used in this experiment are listed in Supplementary Table 4.

### TurboID-based proximity labelling (PL)

TurboID-based PL was performed as described by Zhang *et al*. (2019). Prior to the preparation of samples for MS, the biotin ligase activity of TurboID-containing constructs was tested. In brief, *A. tumefaciens* cells harboring the constructs to express TurboID-fused protein of interest and control constructs were infiltrated into fully expanded leaves of 3- to 4-week-old *N. benthamiana* plants using a 1 mL needleless syringe. Biotin solution (50 μM biotin and 2.5 mM MgCl_2_ in DMSO) or control solution (2.5 mM MgCl_2_ in DMSO) was infiltrated into the same leaf tissues at 42 h post-infiltration. Samples were harvested 6 h after biotin supplementation. Immunoblot assays were performed to detect the protein accumulation and protein biotinylation activity with anti-HA and streptavidin-HRP antibodies, respectively.

Sample preparation for MS was performed as described by Zhang *et al*. (2019) with minor modifications. Briefly, biotin-supplemented samples were harvested as described above and ground to a fine powder in liquid nitrogen, and protein extraction (as described above) and removal of free biotin were carried out. A 100 µL-aliquot of lysate was taken from each sample before and after removal of free biotin. The lysates were incubated with streptavidin-C1-conjugated magnetic beads (Invitrogen, 65001) at 4°C for 3-4 h on a rotator to enrich for biotinylated proteins. In the final wash step, 10% of the suspension was taken to confirm the successful enrichment of biotinylated proteins. Immunoblot analysis of protein enrichment and biotinylation was performed on all lysates taken from the different processing steps.

The antibodies used for TurboID-based PL were Streptavidin-HRP (Abcam, Ab7403; dilution ratio: 1:15,000), anti-HA (Roche, 12013819001; dilution ratio: 1: 5000), and anti-Rat IgG (Abcam, Ab7097; dilution ratio: 1:15,000).

### Statistical analyses

For qPCR and qRT-PCR analyses, the statistical differences between two samples were evaluated using Student’s t-test, with a significance level (α) set at 0.05. To compare the statistical differences between more than two samples, one-way ANOVA followed by Dunnett’s post hoc test was used. Asterisks are used to indicate significant statistical difference (****, P<0.0001; ***, P<0.001; **, P<0.01; *, P<0.05; not significant (ns), P>0.05).

## ACKNOWLEDGEMENTS

The authors thank Axel Giudicatti, Huang Tan, Man Gao, Nicholay Diaz-Ardila, Shuyi Luo, and Zhihao Jiang for critical reading of the manuscript, and Bettina Stadelhofer and the central facilities at the ZMBP, especially the Plant cultivation and the Microscopy facilities, for excellent technical support. This work was partially funded by the European Research Council (GemOmics; 101044142), the Excellence Strategy of the German Federal and State Governments, and the German Research Foundation (DFG) and the National Natural Science Foundation of China (NSFC) through the DFG/NSFC collaboration project 31961133023.

## SUPPLEMENTARY MATERIAL

### SUPPLEMENTARY FIGURES

Supplementary figures 1-21

### SUPPLEMENTARY TABLES

Supplementary tables 1-4

## REFERENCES

Bagewadi B, Chen S, Lal SK, Choudhury NR, Mukherjee SK. 2004. PCNA Interacts with Indian Mung Bean Yellow Mosaic Virus Rep and Downregulates Rep Activity. Journal of Virology 78, 11890–11903.

Bonnamy M, Blanc S, Michalakis Y. 2023. Replication mechanisms of circular ssDNA plant viruses and their potential implication in viral gene expression regulation. (VR Prasad, Ed.). mBio 14, e01692–23.

Castillo AG, Collinet D, Deret S, Kashoggi A, Bejarano ER. 2003. Dual interaction of plant PCNA with geminivirus replication accessory protein (Ren) and viral replication protein (Rep). Virology 312, 381–394.

Choudhury NR, Malik PS, Singh DK, Islam MN, Kaliappan K, Mukherjee SK. 2006. The oligomeric Rep protein of Mungbean yellow mosaic India virus (MYMIV) is a likely replicative helicase. Nucleic Acids Research 34, 6362–6377.

Clérot D, Bernardi F. 2006. DNA Helicase Activity Is Associated with the Replication Initiator Protein Rep of Tomato Yellow Leaf Curl Geminivirus. Journal of Virology 80, 11322–11330.

Cornet L, Shan-e-Ali Z, Li J, Ngapout Y, Shakir S, Meunier L, Callot C, Marande W, Hanikenne M, Rombauts S, Van de Peer Y, Vanderschuren H. 2025. A BAC-guided haplotype assembly pipeline increases the resolution of the virus resistance locus CMD2 in cassava. Genome Biology 26, 185.

Da Silva JPH, De Resende FMP, Da Silva JCF, De Breuil S, Nome C, Bejerman N, Zerbini FM. 2023. Amesuviridae: a new family of plant-infecting viruses in the phylum Cressdnaviricota, realm Monodnaviria. Archives of Virology 168, 223.

Fuchs J, Cheblal A, Gasser SM. 2021. Underappreciated Roles of DNA Polymerase δ in Replication Stress Survival. Trends in Genetics 37, 476–487.

Gassmann M, Focher F, Buhk H-J, Ferrari E, Spadari S, Hübscher U. 1988. Replication of single-stranded porcine circovirus DNA by DNA polymerases α and δ. Biochimica et Biophysica Acta (BBA) - Gene Structure and Expression 951, 280–289.

George B, Ruhel R, Mazumder M, Sharma VK, Jain SK, Gourinath S, Chakraborty S. 2014. Mutational analysis of the helicase domain of a replication initiator protein reveals critical roles of Lys 272 of the B′ motif and Lys 289 of the β-hairpin loop in geminivirus replication. Journal of General Virology 95, 1591–1602.

Gilbert KB, Gallardo P, King SF, Morris CM, Hernandez GL, Carrington JC, Bart RS. 2025. Allelic differences within DNA polymerase delta subunit 1 correlate with geminivirus resistance in diverse plants. (S Smith, Ed.). G3: Genes, Genomes, Genetics 15, jkaf216.

Gronenborn B, Randles J, Vetten H, Thomas J. 2021. Create one new family (Metaxyviridae) with one new genus (Cofodevirus) and one species (Coconut foliar decay virus) moved from the family Nanoviridae (Mulpavirales). International Committee for Taxonomy of Viruses proposal (Taxoprop) number 2020.022P in press.

Hanley-Bowdoin L, Bejarano ER, Robertson D, Mansoor S. 2013. Geminiviruses: masters at redirecting and reprogramming plant processes. Nature Reviews Microbiology 11, 777–788.

He Y-Z, Wang Y-M, Yin T-Y, Fiallo-Olivé E, Liu Y-Q, Hanley-Bowdoin L, Wang X-W. 2020. A plant DNA virus replicates in the salivary glands of its insect vector via recruitment of host DNA synthesis machinery. Proceedings of the National Academy of Sciences 117, 16928–16937.

Hicks WM, Kim M, Haber JE. 2010. Increased Mutagenesis and Unique Mutation Signature Associated with Mitotic Gene Conversion. Science 329, 82–85.

Jeske H. 2009. Geminiviruses. In: De Villiers E-M, Hausen HZ, eds. Current Topics in Microbiology and Immunology. TT Viruses. Berlin, Heidelberg: Springer Berlin Heidelberg, 185–226.

Krupovic M, Varsani A, Kazlauskas D, et al. 2020. *Cressdnaviricota* : a Virus Phylum Unifying Seven Families of Rep-Encoding Viruses with Single-Stranded, Circular DNA Genomes. (RM Sandri-Goldin, Ed.). Journal of Virology 94, e00582–20.

Lal A, Vo TTB, Sanjaya IGNPW, Ho PT, Kim J-K, Kil E-J, Lee S. 2020. Nanovirus Disease Complexes: An Emerging Threat in the Modern Era. Frontiers in Plant Science 11, 558403.

Langston LD, O’Donnell M. 2008. DNA Polymerase δ Is Highly Processive with Proliferating Cell Nuclear Antigen and Undergoes Collision Release upon Completing DNA. Journal of Biological Chemistry 283, 29522–29531.

Laufs J, Traut W, Heyraud F, Matzeit V, Rogers SG, Schell J, Gronenborn B. 1995. In vitro cleavage and joining at the viral origin of replication by the replication initiator protein of tomato yellow leaf curl virus. Proceedings of the National Academy of Sciences 92, 3879–3883.

Li F, Xu X, Li Z, Wang Y, Zhou X. 2020. Identification of Yeast Factors Involved in the Replication of Mungbean Yellow Mosaic India Virus Using Yeast Temperature-Sensitive Mutants. Virologica Sinica 35, 120–123.

Lim Y-W, Mansfeld BN, Schläpfer P, et al. 2022. Mutations in DNA polymerase δ subunit 1 co-segregate with CMD2-type resistance to Cassava Mosaic Geminiviruses. Nature Communications 13, 3933.

Luque A, Sanz-Burgos AP, Ramirez-Parra E, Castellano MM, Gutierrez C. 2002a. Interaction of Geminivirus Rep Protein with Replication Factor C and Its Potential Role during Geminivirus DNA Replication. Virology 302, 83–94.

Luque A, Sanz-Burgos AP, Ramirez-Parra E, Castellano MM, Gutierrez C. 2002b. Interaction of Geminivirus Rep Protein with Replication Factor C and Its Potential Role during Geminivirus DNA Replication. Virology 302, 83–94.

Maimbo M, Ohnishi K, Hikichi Y, Yoshioka H, Kiba A. 2010. S-Glycoprotein-Like Protein Regulates Defense Responses in *Nicotiana* Plants against *Ralstonia solanacearum*. Plant Physiology 152, 2023–2035.

Maio F, Arroyo-Mateos M, Bobay BG, Bejarano ER, Prins M, Van Den Burg HA. 2019. A Lysine Residue Essential for Geminivirus Replication Also Controls Nuclear Localization of the Tomato Yellow Leaf Curl Virus Rep Protein. (AE Simon, Ed.). Journal of Virology 93, e01910–18.

Mason G, Caciagli P, Accotto GP, Noris E. 2008. Real-time PCR for the quantitation of Tomato yellow leaf curl Sardinia virus in tomato plants and in Bemisia tabaci. Journal of Virological Methods 147, 282–289.

Nakagawa T, Kurose T, Hino T, Tanaka K, Kawamukai M, Niwa Y, Toyooka K, Matsuoka K, Jinbo T, Kimura T. 2007a. Development of series of gateway binary vectors, pGWBs, for realizing efficient construction of fusion genes for plant transformation. Journal of Bioscience and Bioengineering 104, 34–41.

Nakagawa T, Suzuki T, Murata S, et al. 2007b. Improved Gateway Binary Vectors: High-Performance Vectors for Creation of Fusion Constructs in Transgenic Analysis of Plants. Bioscience, Biotechnology, and Biochemistry 71, 2095–2100.

Nash K, Chen W, Muzyczka N. 2008. Complete In Vitro Reconstitution of Adeno-Associated Virus DNA Replication Requires the Minichromosome Maintenance Complex Proteins. Journal of Virology 82, 1458–1464.

Pott DM, Gao M, Shi C, et al. 2025. Pervasive splicing in a plant DNA virus. Plant Biology doi: 10.1101/2025.10.01.679800. [Preprint].

Pott DM, Medina-Puche L, Shi C, et al. 2026. Alternative splicing expands the functional portfolio of a plant virus to control the viral cycle. Plant Biology doi: 10.64898/2026.02.19.706752. [Preprint].

Preiss W, Jeske H. 2003. Multitasking in Replication Is Common among Geminiviruses. Journal of Virology 77, 2972–2980.

Rojas MR, Macedo MA, Maliano MR, et al. 2018. World Management of Geminiviruses. Annual Review of Phytopathology 56, 637–677.

Ruhel R, Mazumder M, Gnanasekaran P, Kumar M, Gourinath S, Chakraborty S. 2021. Functional implications of residues of the B′ motif of geminivirus replication initiator protein in its helicase activity. The FEBS Journal 288, 6492–6509.

Shen X, Gill U, Arens M, Yan Z, Bai Y, Hutton SF, Wolters A-MA. 2025. The tomato gene Ty-6, encoding DNA polymerase delta subunit 1, confers broad resistance to Geminiviruses. Theoretical and Applied Genetics 138, 22.

Singh DK, Islam MN, Choudhury NR, Karjee S, Mukherjee SK. 2007. The 32 kDa subunit of replication protein A (RPA) participates in the DNA replication of Mung bean yellow mosaic India virus (MYMIV) by interacting with the viral Rep protein. Nucleic Acids Research 35, 755–770.

Siskos L, Antoniou M, Riado J, et al. 2023. DNA primase large subunit is an essential plant gene for geminiviruses, putatively priming viral ss-DNA replication. Frontiers in Plant Science 14, 1130723.

Timchenko T, De Kouchkovsky F, Katul L, David C, Vetten HJ, Gronenborn B. 1999. A Single Rep Protein Initiates Replication of Multiple Genome Components of Faba Bean Necrotic Yellows Virus, a Single-Stranded DNA Virus of Plants. Journal of Virology 73, 10173–10182.

Tsurimoto T, Stillman B. 1991a. Replication factors required for SV40 DNA replication in vitro. I. DNA structure-specific recognition of a primer-template junction by eukaryotic DNA polymerases and their accessory proteins. Journal of Biological Chemistry 266, 1950–1960.

Tsurimoto T, Stillman B. 1991b. Replication factors required for SV40 DNA replication in vitro. II. Switching of DNA polymerase alpha and delta during initiation of leading and lagging strand synthesis. Journal of Biological Chemistry 266, 1961–1968.

Varma A, Malathi VG. 2003. Emerging geminivirus problems: A serious threat to crop production. Annals of Applied Biology 142, 145–164.

Wang L, Tan H, Medina-Puche L, et al. 2022. Combinatorial interactions between viral proteins expand the potential functional landscape of the tomato yellow leaf curl virus proteome. (DM Bisaro, Ed.). PLOS Pathogens 18, e1010909.

Wu M, Wei H, Tan H, Pan S, Liu Q, Bejarano ER, Lozano-Durán R. 2021. Plant DNA polymerases α and δ mediate replication of geminiviruses. Nature Communications 12, 2780.

Zhao L, Rosario K, Breitbart M, Duffy S. 2019. Eukaryotic Circular Rep-Encoding Single-Stranded DNA (CRESS DNA) Viruses: Ubiquitous Viruses With Small Genomes and a Diverse Host Range. Advances in Virus Research. Elsevier, 71–133.

