## Supplementary material for "Selective repurposing of the eukaryotic DNA replication machinery by a plant virus"

### SUPPLEMENTARY FIGURES

Supplementary figures 1-21

### SUPPLEMENTARY TABLES

Supplementary tables 1-5

### SUPPLEMENTARY FIGURES

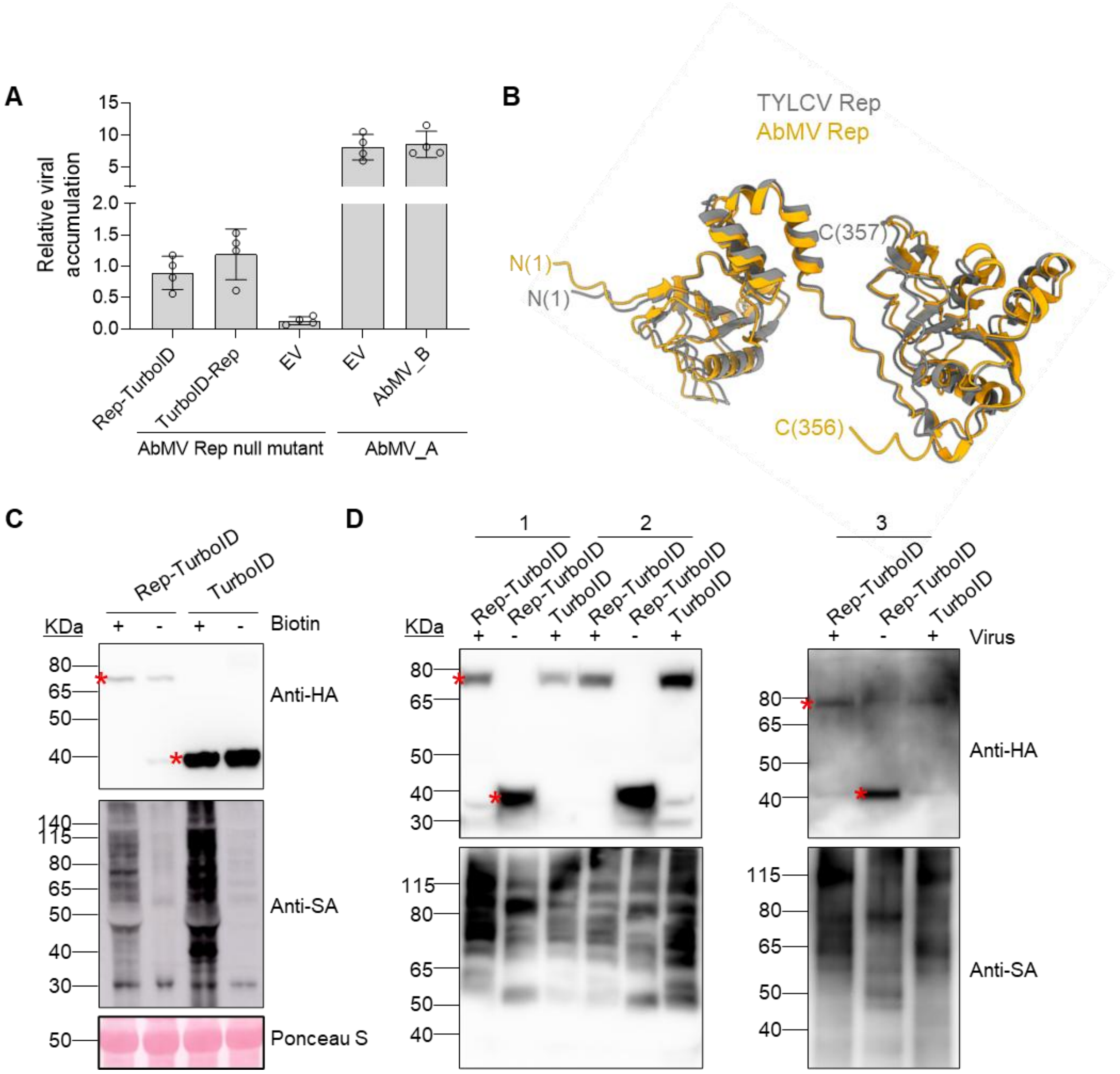


**Figure S1. AbMV Rep fused to TurboID either N- or C-terminally can mediate viral genome replication and retains biotin ligase activity. A.** Rep from AbMV fused to either N- and C-terminal TurboID tag can mediate the replication of viral genome. A AbMV Rep null mutant infectious clone was transiently co-transformed with constructs to express Rep fusion proteins or an empty vector (EV; negative control) in *N. benthamiana* leaves. Co-transformation of AbMV A component with EV or AbMV component B were used as the positive controls. Samples were harvested at 3 days post-infiltration. Viral DNA accumulation was measured by qPCR with 25S ribosomal DNA interspacer (ITS) as the internal reference. This experiment was repeated three times with similar results. **B.** Structural overlay of AbMV Rep onto TYLCV Rep, as predicted by AlphaFold 3. TYLCV Rep and AbMV Rep are shown in grey and orange, respectively. **C.** Immunoblot analysis of protein accumulation (anti-HA) and biotinylation activity (Streptavidin-HRP, anti-SA) in the presence and absence of exogenous biotin. At 42 hours post-infiltration, *N. benthamiana* leaves were infiltrated with either 50 μM biotin (+Biotin) or DMSO (-Biotin) and harvested 6 hours later. Ponceau S, ponceau staining. **D.** Immunoblot analysis of fusion protein level (anti-HA) and biotinylated proteins (anti-SA) in the samples from input lysates (1), after desalting (2), and after affinity purification using streptavidin beads (3). Virus, AbMV Rep null mutant. This experiment was repeated three times with similar results. The position of asterisks in panels C and D indicate the expected bands of Rep-TurboID (~78.0 kDa) and TurboID (~38.7 kDa), respectively.

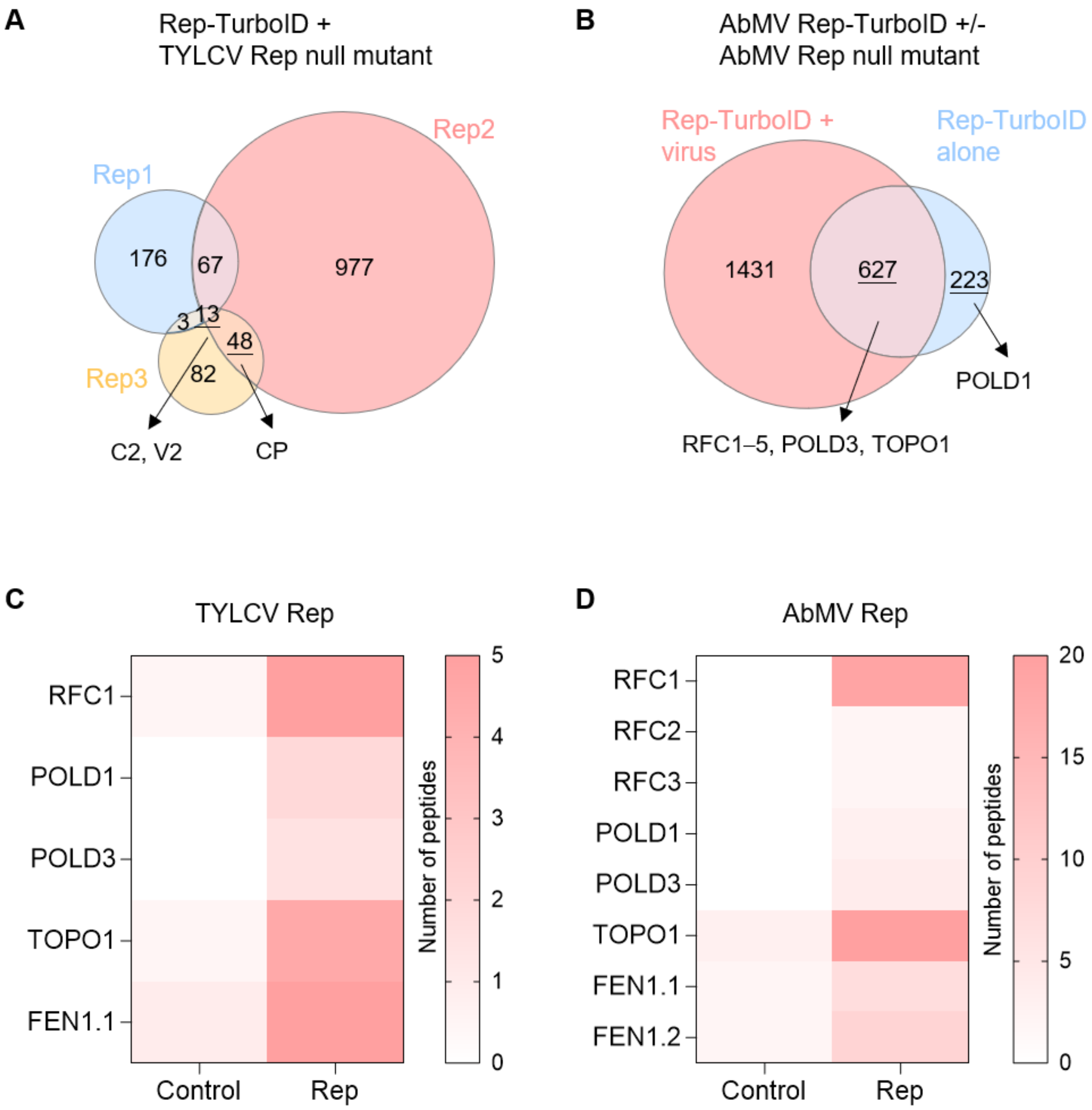


**Figure S2. Proteins commonly labelled by TYLCV Rep and AbMV Rep. A.** Venn diagram showing the overlap of TYLCV Rep-labelled proteins across three independent biological replicates (Rep1-3) in the presence of virus infection. **B.** Venn diagram showing the overlap of Rep-labelled proteins in the presence and absence of viral infection for AbMV. Arrows indicate DNA replication-related proteins specifically labelled under different experimental conditions. **C-D.** Heatmap depicting the abundance of peptides in the presence (Rep) and absence (control) of Rep across biological replicates after filtering, for TYLCV Rep (C) and AbMV Rep (D).

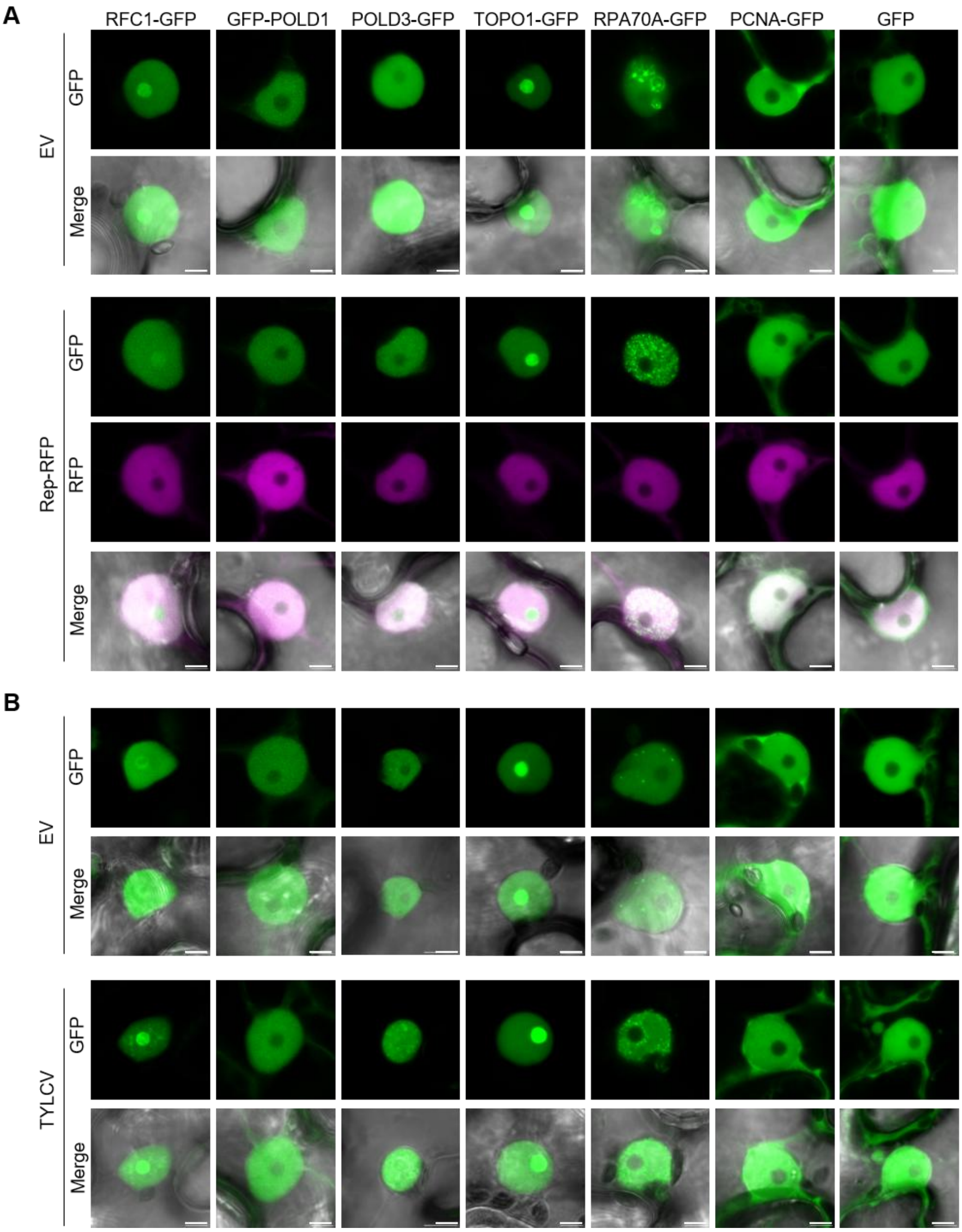


**Figure S3. Subcellular localization of selected DNA replication-related proteins. A-B.** Transient expression of RFC1-GFP, GFP-POLD1, POLD3-GFP, TOPO1-GFP, RPA70A-GFP, PCNA-GFP, and GFP in *N. benthamiana* leaves, either in the absence (EV, co-transformed with an empty vector) or presence of Rep-RFP (A) or TYLCV (B). *Agrobacterium* cells containing the respective binary vectors were co-infiltrated at a 1:1 ratio; GFP was used as a negative control; confocal images were taken at 30 hours post-infiltration. Merge, overlay of fluorescence and bright-field images. Scale bar: 5 μm. This experiment was repeated twice with similar results.

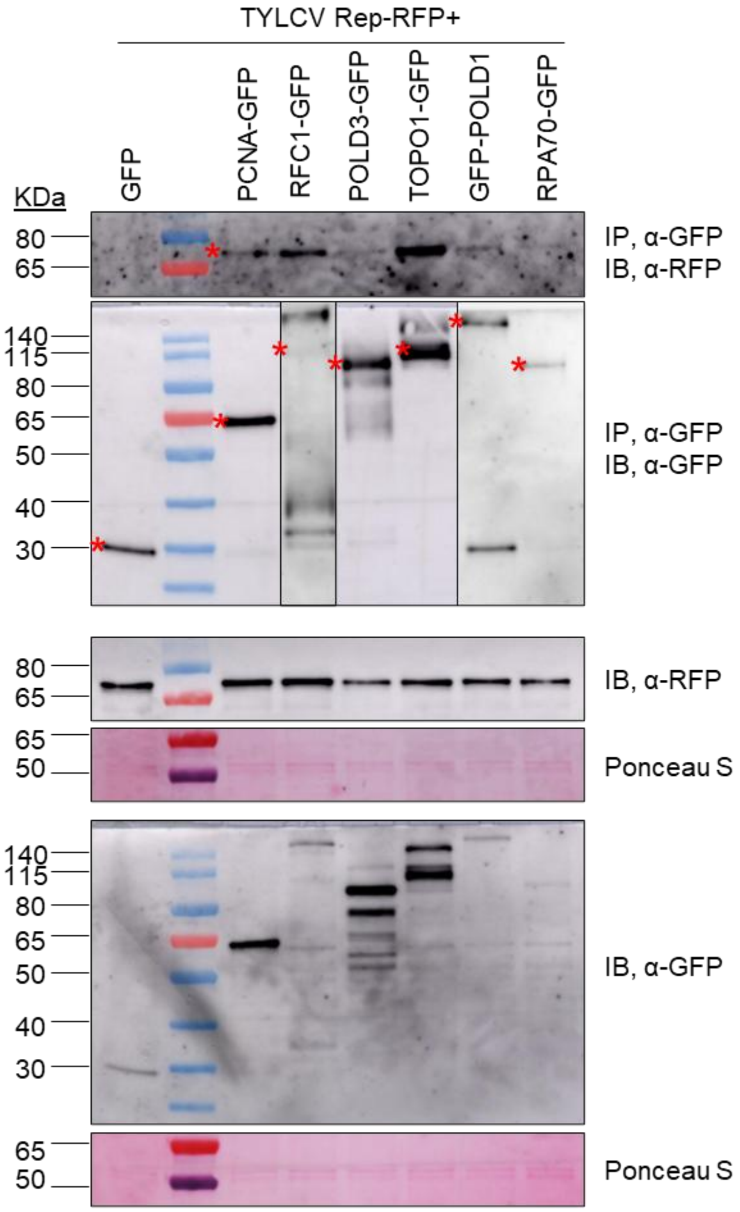


**Figure S4. Rep associates with selected replication-related factors in co-immunoprecipitation experiments.** Rep-RFP co-immunoprecipitates with RFC1-GFP, POLD3-GFP, TOPO1-GFP, GFP-POLD1, RPA70A-GFP, and PCNA-GFP (positive control), but not with GFP (negative control), upon transient expression in *N. benthamiana*. IP, immunoprecipitate; IB, immunoblotting; Ponceau S, ponceau staining. The position of asterisks indicates the expected bands: RFC1-GFP, ~134 kDa; POLD3-GFP, ~85 kDa; TOPO1-GFP, ~125 kDa; GFP-POLD1, ~148 kDa; RPA70A-GFP, ~99 kDa; PCNA-GFP, ~58 kDa; GFP, ~28 kDa. This experiment was repeated three times with similar results.

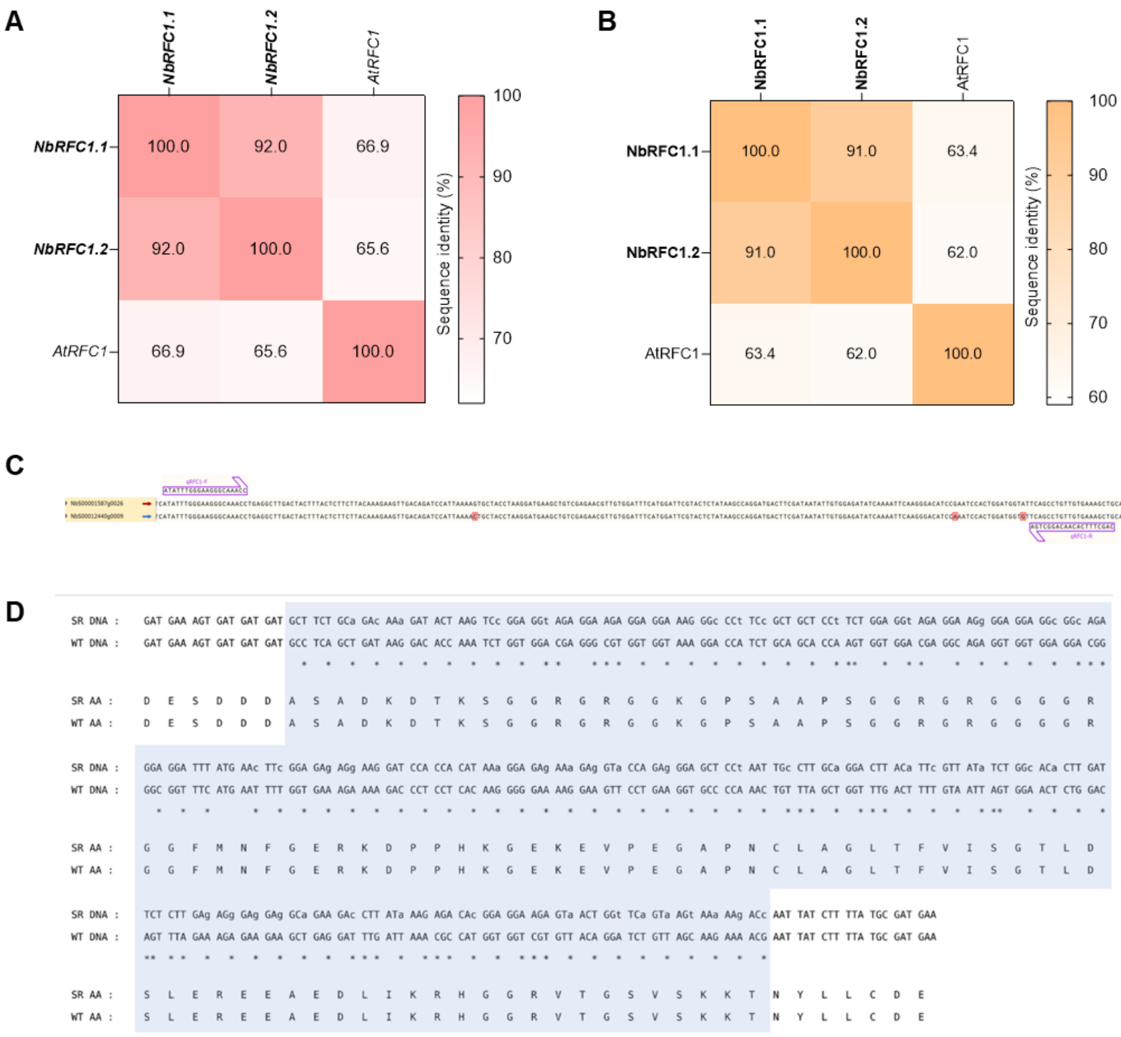

**Figure S5. Sequence analysis of *N. benthamiana RFC1* homologs and RFC1 isoforms, qPCR primer positions for expression quantification, and design of silencing-resistant coding sequence variants. A-B.** Pairwise sequence identity analysis of RFC1 isoforms/*RFC1* homologs and their Arabidopsis ortholog. DNA (A) and protein (B) sequence identity matrices. NbRFC1.1 (NbS00001587g0026), NbRFC1.2 (NbS00012440g0009), and AtRFC1 (AT5G22010) are shown in the figure. Values represent pairwise sequence identities (%). Isoforms silenced in this study are shown in bold. **C.** Alignment of isoform 1-derived qPCR primers with other *N. benthamiana NbRFC1* isoforms. **D.** Alignment of silencing-resistant (SR) and wild-type (WT) coding sequences and the corresponding resulting protein. The VIGS-targeted region (~300 bp) is highlighted in grey. Synonymous nucleotide substitutions were introduced into the SR sequence within this region while preserving the encoded amino acid sequence. Only short flanking sequences are shown for context.

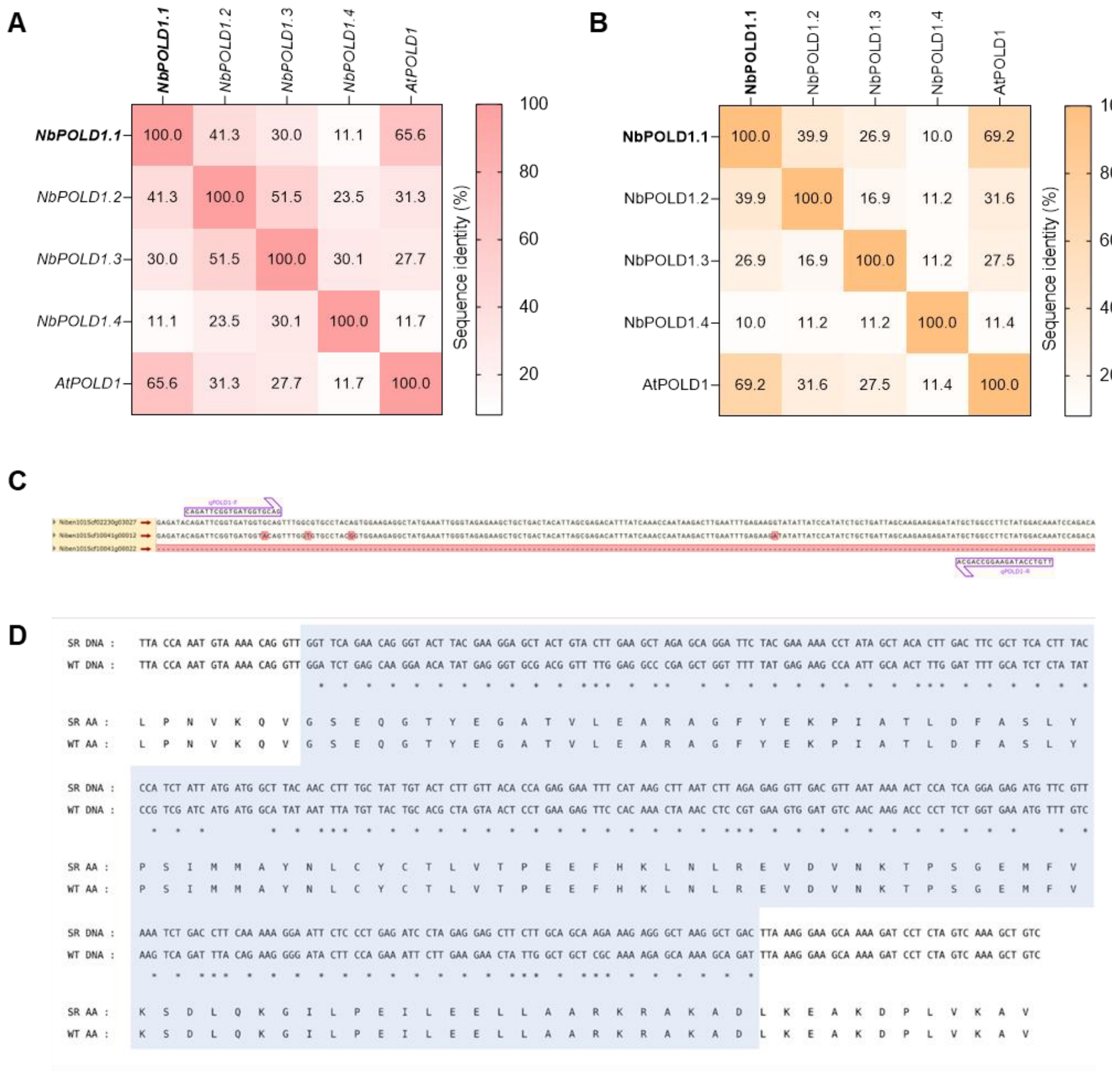


**Figure S6. Sequence analysis of *N. benthamiana POLD1* homologs and POLD1 isoforms, qPCR primer positions for expression quantification, and design of silencing-resistant coding sequence variants. A-B.** Pairwise sequence identity analysis of POLD1 isoforms/*POLD1* homologs and their Arabidopsis ortholog. DNA (A) and protein (B) sequence identity matrices. NbPOLD1.1 (Niben101Scf02230g03027), NbPOLD1.2 (Niben101Scf00215g00022), NbPOLD1.3 (Niben101Scf10041g00012), NbPOLD1.4 (Niben101Scf10041g00022) and AtPOLD1 (AT5G63960) are shown in the figure. Values represent pairwise sequence identities (%). Isoforms silenced in this study are shown in bold. **C.** Alignment of isoform 1-derived qPCR primers with other *N. benthamiana NbPOLD1* isoforms. **D.** Alignment of silencing-resistant (SR) and wild-type (WT) coding sequences and the corresponding resulting protein. The VIGS-targeted region (~300 bp) is highlighted in grey. Synonymous nucleotide substitutions were introduced into the SR sequence within this region while preserving the encoded amino acid sequence. Only short flanking sequences are shown for context.

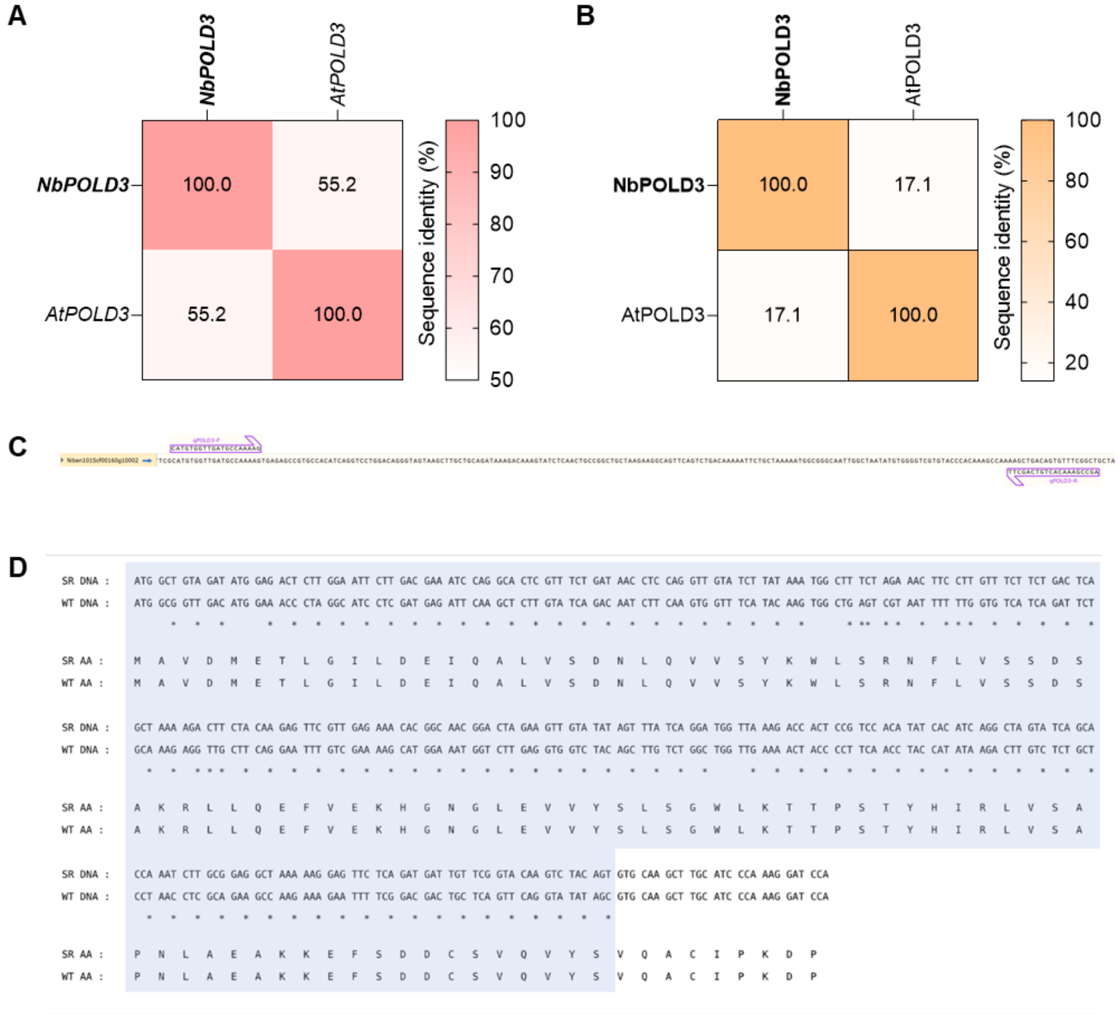


**Figure S7. Sequence analysis of *N. benthamiana POLD3* homologs and POLD3 isoforms, qPCR primer positions for expression quantification, and design of silencing-resistant coding sequence variants. A-B.** Pairwise sequence identity analysis of POLD3 isoform/*POLD3* homolog and its Arabidopsis ortholog. DNA (A) and protein (B) sequence identity matrices. NbPOLD3 (Niben101Scf00160g10002) and AtPOLD3 (AT1G78650) are shown in the figure. Values represent pairwise sequence identities (%). Isoform silenced in this study is shown in bold. **C.** Alignment of isoform 1-derived qPCR primers with other *N. benthamiana NbPOLD3* isoforms. **D.** Alignment of silencing-resistant (SR) and wild-type (WT) coding sequences and the corresponding resulting protein. The VIGS-targeted region (~300 bp) is highlighted in grey. Synonymous nucleotide substitutions were introduced into the SR sequence within this region while preserving the encoded amino acid sequence. Only short flanking sequences are shown for context.

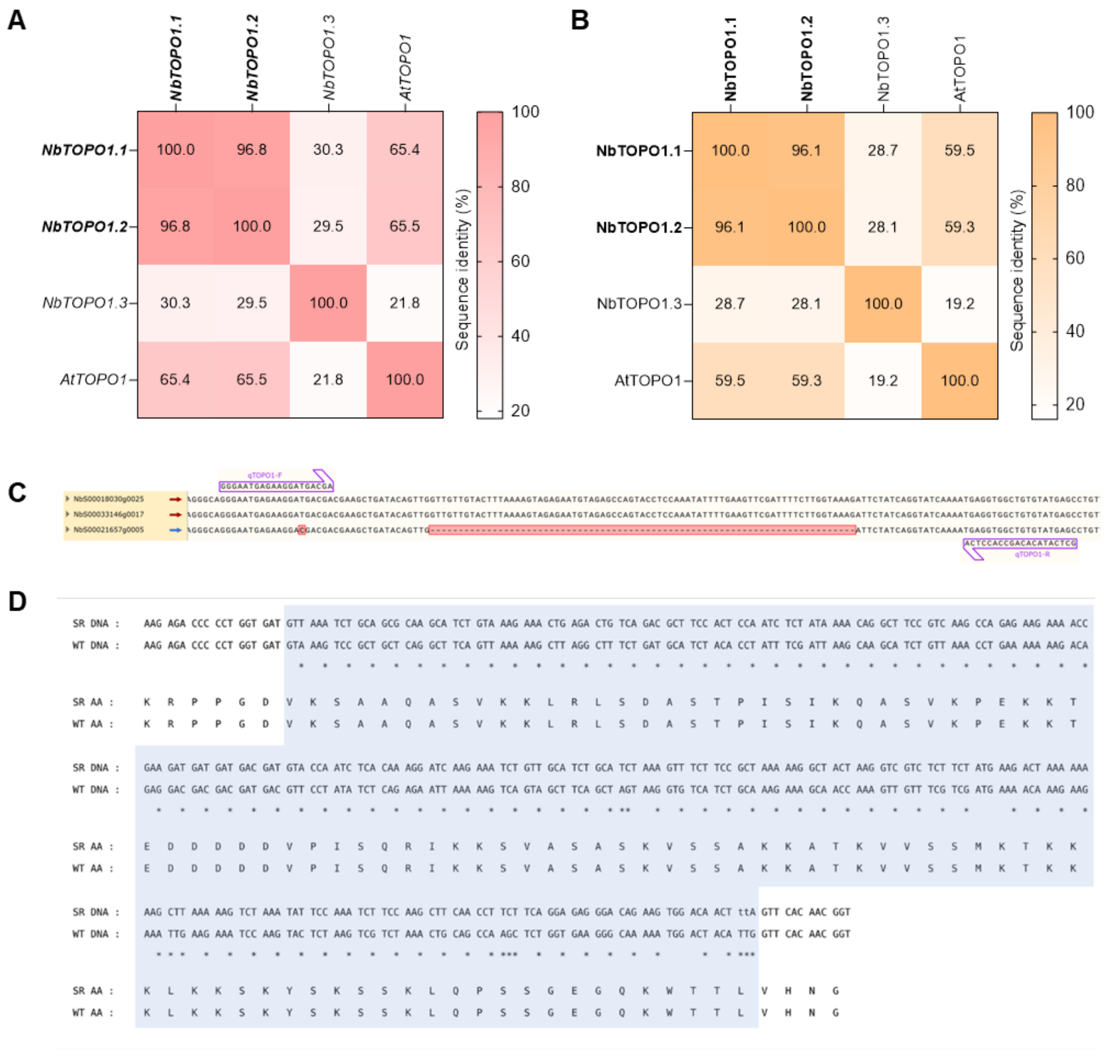


**Figure S8. Sequence analysis of *N. benthamiana TOPO1* homologs and TOPO1 isoforms, qPCR primer positions for expression quantification, and design of silencing-resistant coding sequence variants. A-B.** Pairwise sequence identity analysis of TOPO1 isoforms/*TOPO1* homologs and their Arabidopsis ortholog. DNA (A) and protein (B) sequence identity matrices. NbTOPO1.1 (NbS00018030g0025), NbTOPO1.2 (NbS00033146g0017), NbTOPO1.3 (NbS00021657g0005), and AtTOPO1 (AT5G55310) are shown in the figure. Values represent pairwise sequence identities (%). Isoforms silenced in this study are shown in bold. **C.** Alignment of isoform 1-derived qPCR primers with other *N. benthamiana NbTOPO1* isoforms. **D.** Alignment of silencing-resistant (SR) and wild-type (WT) coding sequences and the corresponding resulting protein. The VIGS-targeted region (~300 bp) is highlighted in grey. Synonymous nucleotide substitutions were introduced into the SR sequence within this region while preserving the encoded amino acid sequence. Only short flanking sequences are shown for context.

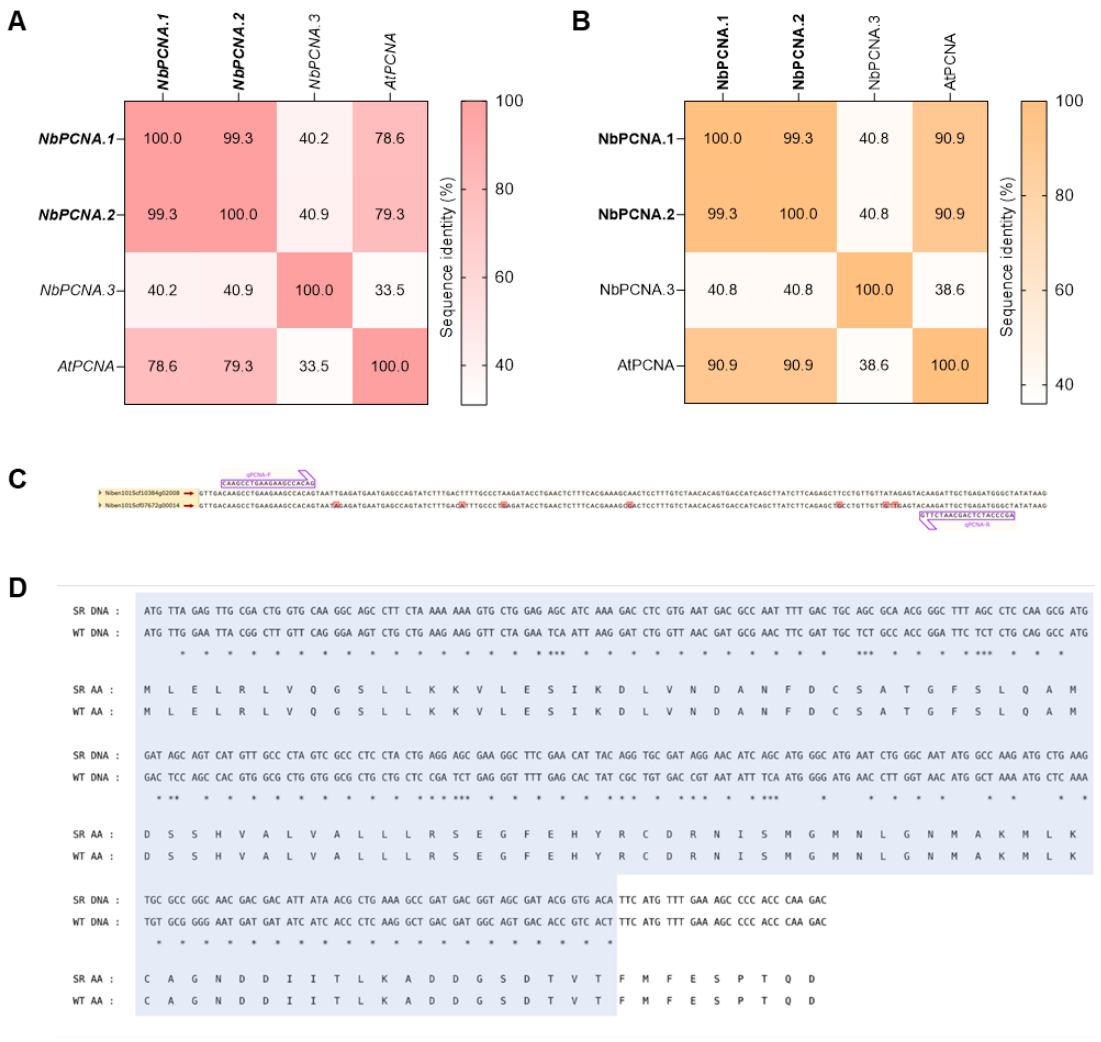


**Figure S9. Sequence analysis of *N. benthamiana PCNA* homologs and PCNA isoforms, qPCR primer positions for expression quantification, and design of silencing-resistant coding sequence variants. A-B.** Pairwise sequence identity analysis of PCNA isoforms/*PCNA* homologs and their Arabidopsis ortholog. DNA (A) and protein (B) sequence identity matrices. NbPCNA.1 (Niben101Scf10384g02008), NbPCNA.2 (Niben101Scf07672g00014), NbPCNA.3 (Niben101Scf07672g00015), and AtPCNA (AT1G07370) are shown in the figure. Values represent pairwise sequence identities (%). Isoforms silenced in this study are shown in bold. **C.** Alignment of isoform 1-derived qPCR primers with other *N. benthamiana NbPCNA* isoforms. **D.** Alignment of silencing-resistant (SR) and wild-type (WT) coding sequences and the corresponding resulting protein. The VIGS-targeted region (~300 bp) is highlighted in grey. Synonymous nucleotide substitutions were introduced into the SR sequence within this region while preserving the encoded amino acid sequence. Only short flanking sequences are shown for context.

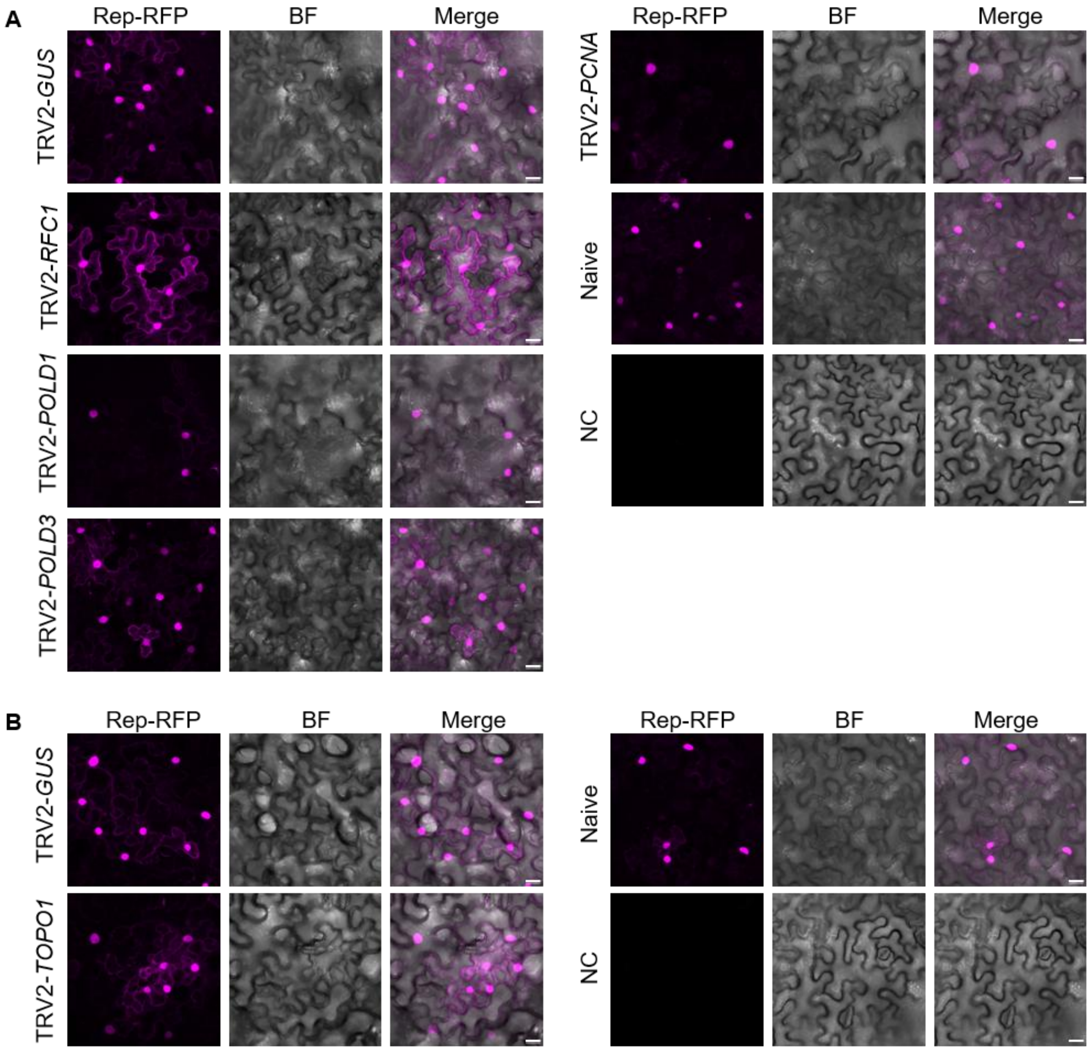


**Figure S10. *Agrobacterium tumefaciens*-mediated transient expression of Rep-RFP is not impaired in *RFC1*-, *POLD1*-, *POLD3*-, *PCNA*- and *TOPO1*-silenced *N. benthamiana* plants.** *Agrobacterium* cells containing TRV1 and TRV2-RFC1, TRV2-POLD1, TRV2-POLD3, TRV2-PCNA, and TRV2-TOPO1 were mixed at a 1:1 ratio and inoculated into 2-week-old *N. benthamiana* seedlings. Plants were then agroinfiltrated with Rep-RFP at 14 days post-TRV inoculation, except for *TOPO1*-silenced plants, which were infiltrated at 7 dpi. Confocal images were taken at 30 hours post-infiltration. BF, brightfield. Merge, overlay of fluorescence and BF images. Scale bar: 20 μm. TRV2-GUS-inoculated and naïve samples served as positive controls, while NC represents the negative control (infiltrated with an empty vector). This experiment was repeated twice with similar results.

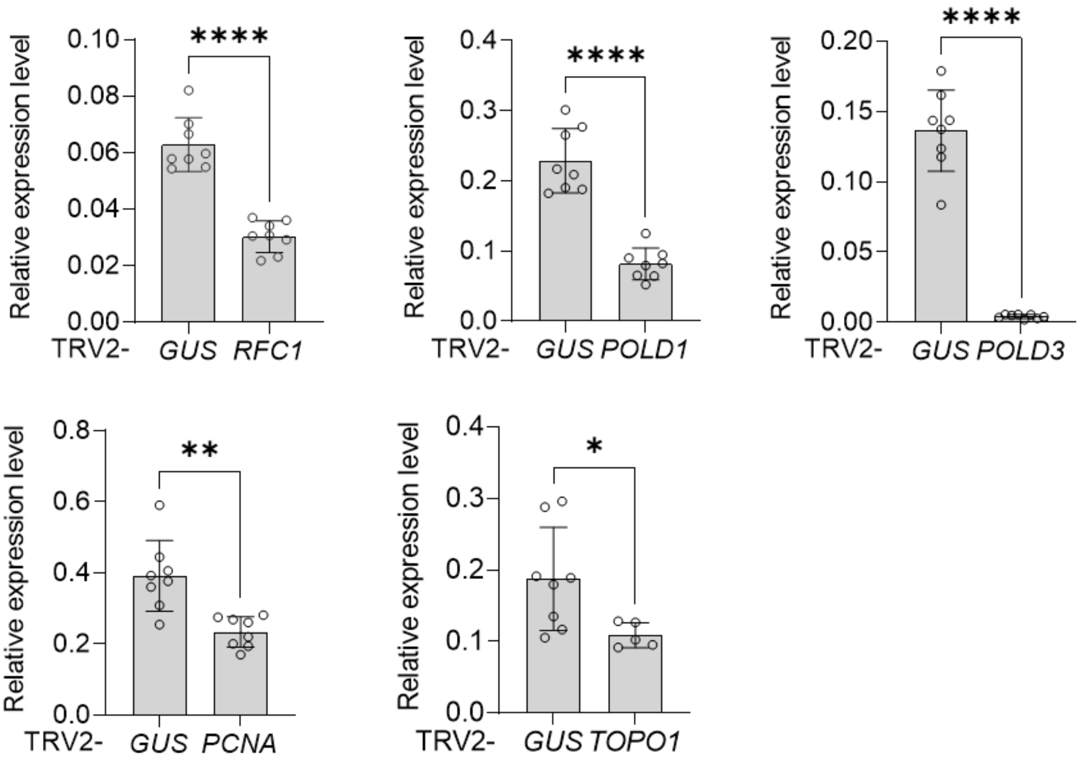


**Figure S11. Silencing efficiency in gene-silenced plants in Figure 2E.** qRT-PCR analysis of silencing efficiency in *RFC1*-, *POLD1*-, *POLD3*-, *PCNA*-, and *TOPO1*-silenced WT *N. benthamiana* plants shown in Figure 2E following agroinfiltration with the TYLCV infectious clone. *NbActin* served as an internal reference gene. Each dot represents an independent plant. Error bars indicate standard deviation. TRV2-GUS plants served as the control. Asterisks indicate statistically significant differences based on Student's t-test (****, P<0.0001; ***, P<0.001; **, P<0.01; *, P<0.05; not significant (ns), P>0.05). All experiments were repeated at least twice with similar results.

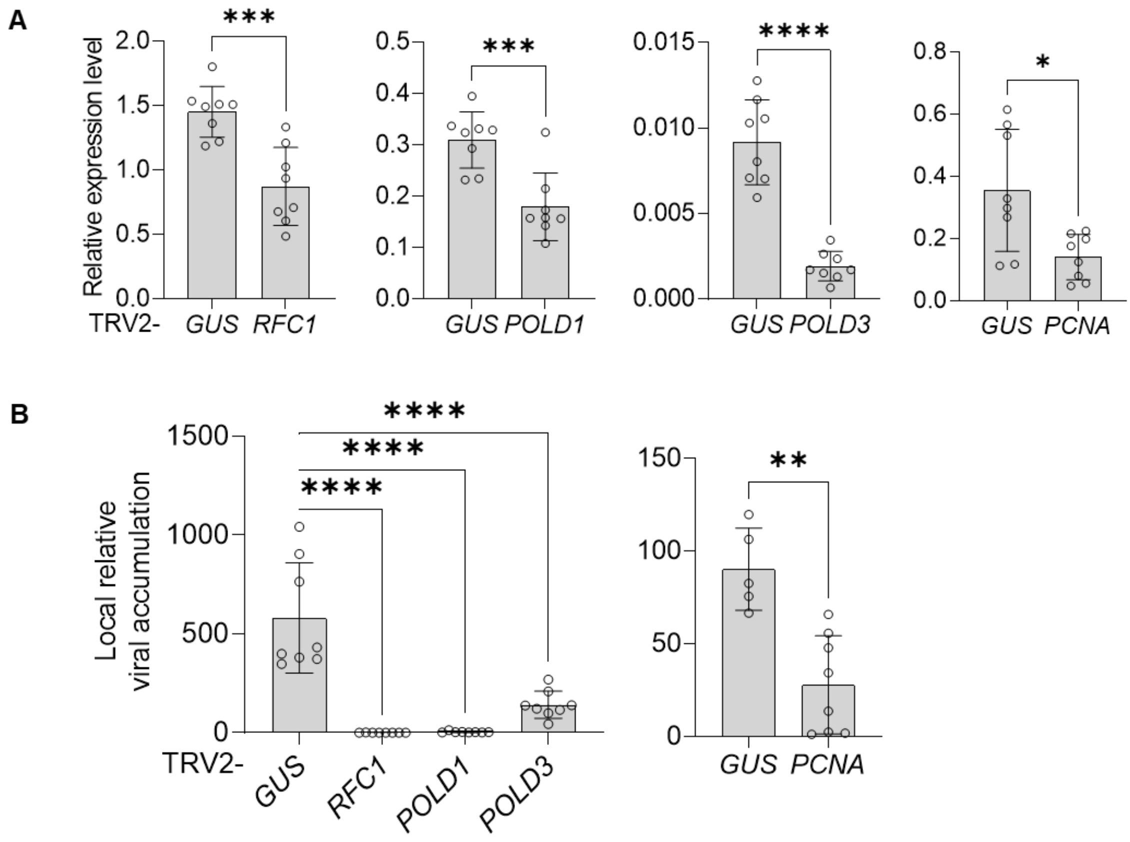


**Figure S12. Selected replication-associated factors are required for replication of the bipartite geminivirus AbMV. A-B.** Silencing efficiency (A) in *RFC1*-, *POLD1*-, *POLD3*-, and *PCNA*-silenced WT *N. benthamiana* plants following agroinfiltration with AbMV A component, measured by qRT-PCR using *NbActin* as an internal reference, and viral accumulation (B), measured by qPCR with 25S ribosomal DNA interspacer (ITS) as an internal reference. All samples were harvested at 3 days post-infiltration. This experiment was repeated twice with similar results. Each dot represents an independent plant. Error bars indicate standard deviation. TRV2-GUS plants served as the control. Asterisks in panel A and on the right side of panel B indicate statistically significant differences based on Student's t-test, whereas asterisks on the left side of panel B indicate statistically significant differences based on one-way ANOVA followed by Dunnett's multiple comparisons test (****, P<0.0001; ***, P<0.001; **, P<0.01; *, P<0.05; not significant (ns), P>0.05).

A

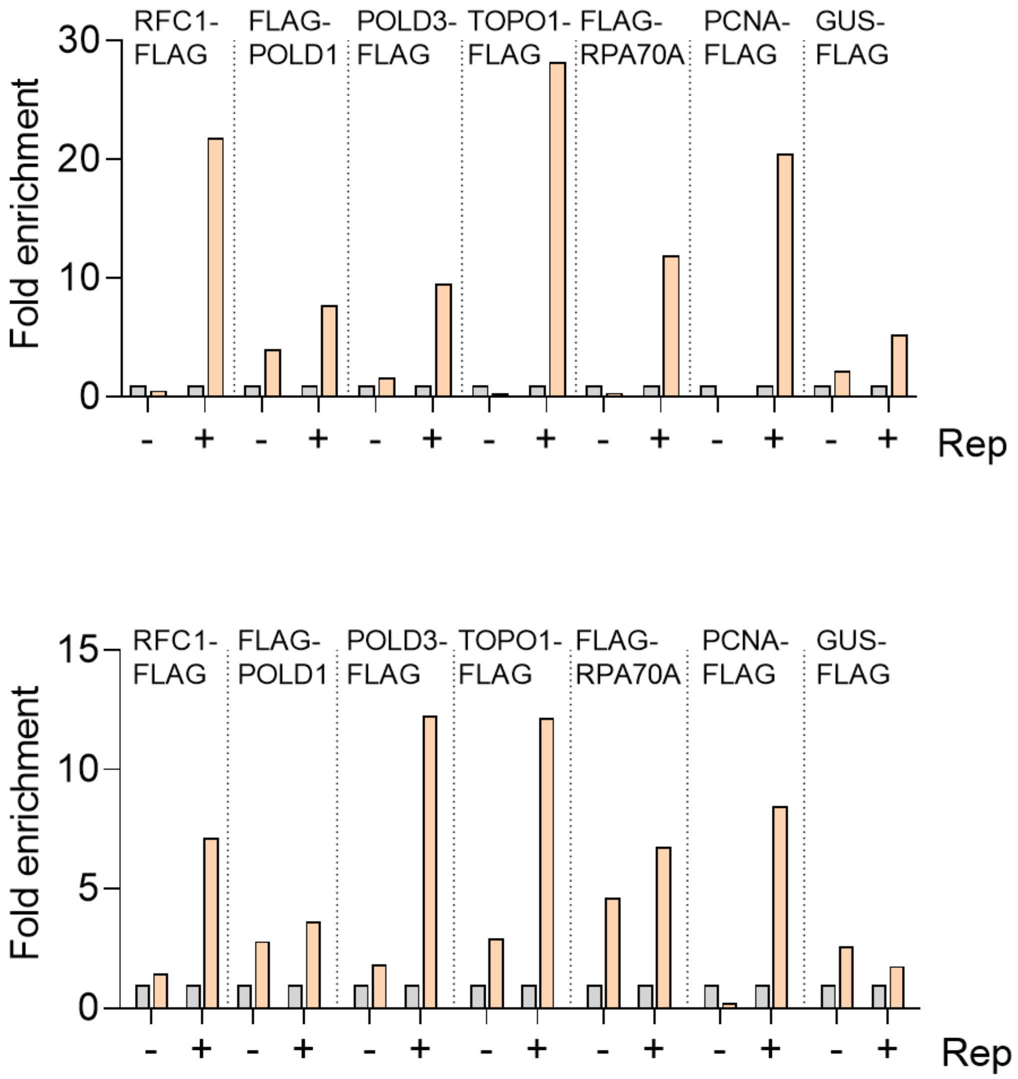


B



**Figure S13. Additional independent biological replicates of the ChIP experiment shown in Figure 4E.** Independent biological replicate 2 (A) and replicate 3 (B). See the legend to Figure 4E for experimental details.

**A**

| Name | Gene identifier | Annotation | Source | EV | TYLCV |
|---|---|---|---|---|---|
| *FEN1.1* | NbS00024431g0002 | Flap endonuclease 1 | TYLCV Rep & AbMV Rep | 202 | 191 |
| *FEN1.2* | NbS00017492g0017 | | AbMV Rep | 147 | 107 |
| *LIG1.1* | Niben101Scf19975g01014 | DNA ligase 1 | Functional correlation | 360 | 503 |
| *LIG1.2* | Niben101Scf00436g03006 | | | 49 | 84 |

**B**

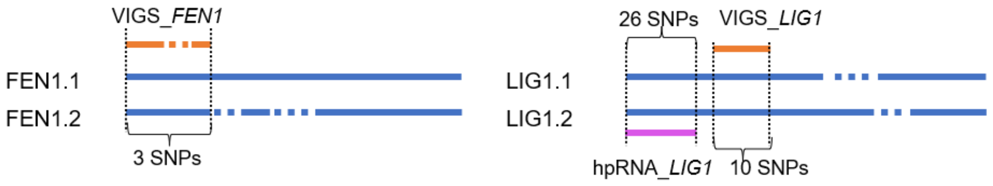


**Figure S14. Expression level and silencing strategy for *FEN1* and *LIG1*. A.** Expression of each orthologue gene in *N. benthamiana* leaves. Expression data are taken from Wu et al. (2019) and represent transcription abundance measured in RPM (reads per million). Data were obtained from leaves transiently transformed with a TYLCV infectious clone or an empty vector (EV) as negative control. **B.** Comparison of each FEN1/LIG1 paralog and the position of the VIGS/hairpin RNA (hpRNA) target sequences. The orange and pink lines represent the VIGS and hpRNA sequence, respectively, while the blue lines represent the gene coding sequence. Dotted lines indicate nucleotide sequences absent from the corresponding paralog. SNPs, single-nucleotide polymorphisms. To simultaneously silence both *LIG1* paralogs, both VIGS- and hpRNA-mediated silencing approaches were used.

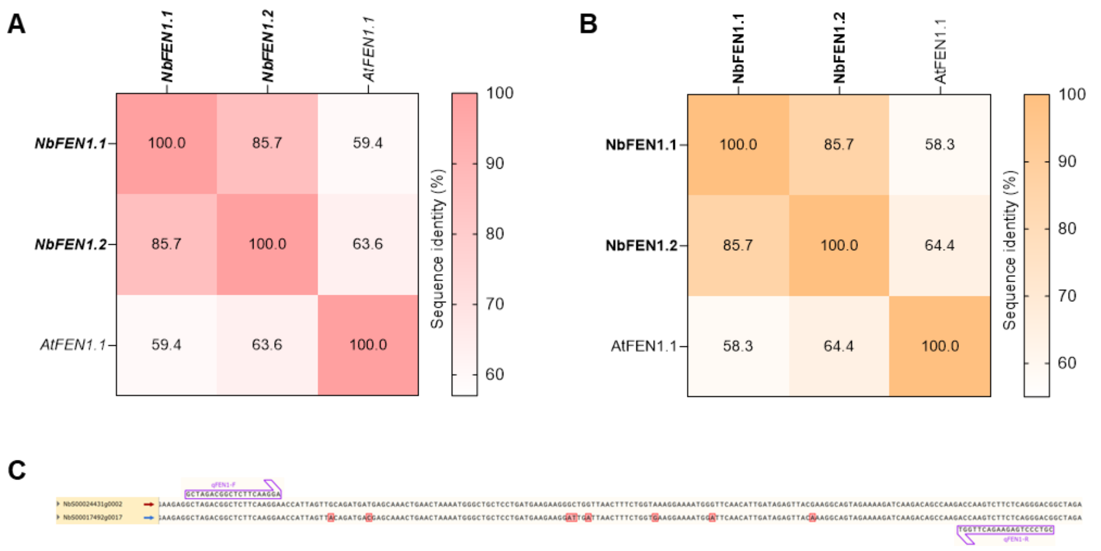


**Figure S15. Sequence analysis of *N. benthamiana FEN1* homologs and FEN1 isoforms, qPCR primer positions for expression quantification. A-B.** Pairwise sequence identity analysis of FEN1 isoforms/*FEN1* homologs and their Arabidopsis ortholog. DNA (A) and protein (B) sequence identity matrices. NbFEN1.1 (NbS00024431g0002), NbFEN1.2 (NbS00017492g0017), and AtFEN1.1 (AT5G26680.1) are shown in the figure. Values represent pairwise sequence identities (%). Isoforms silenced in this study are shown in bold. **C.** Alignment of isoform 1-derived qPCR primers with other *N. benthamiana NbFEN1* isoforms.

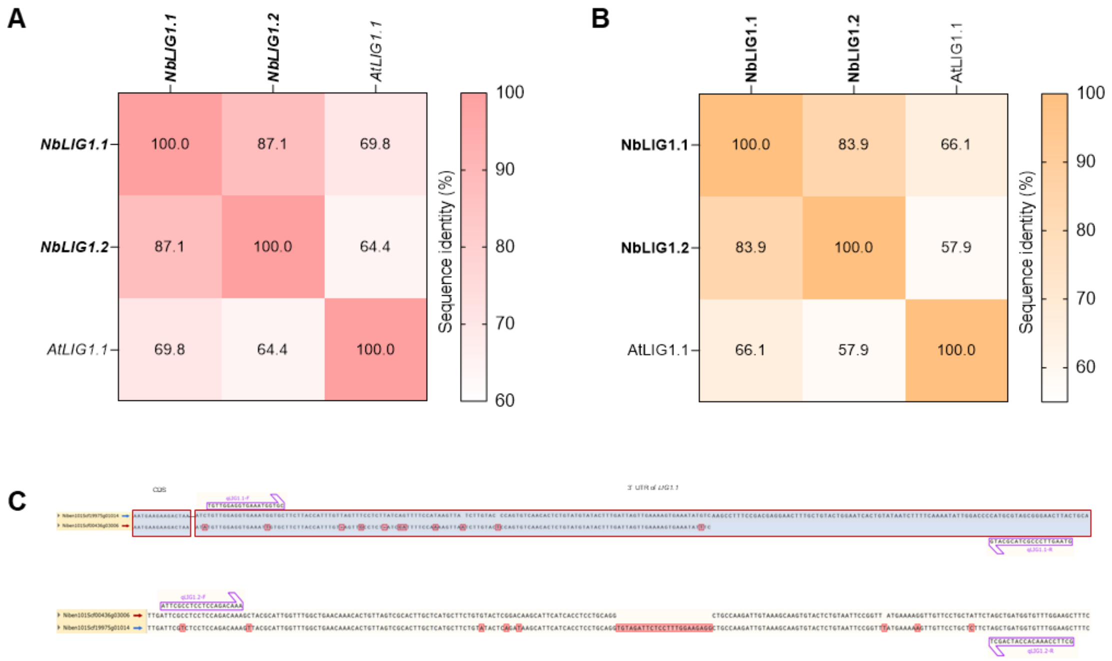


**Figure S16. Sequence analysis of *N. benthamiana LIG1* homologs and LIG1 isoforms, qPCR primer positions for expression quantification. A-B.** Pairwise sequence identity analysis of LIG1 isoforms/*LIG1* homologs and their Arabidopsis ortholog. DNA (A) and protein (B) sequence identity matrices. NbLIG1.1 (Niben101Scf19975g01014), NbLIG1.2 (Niben101Scf00436g03006), and AtLIG1.1 (AT1G08130.1) are shown in the figure. Values represent pairwise sequence identities (%). Isoforms silenced in this study are shown in bold. **C.** Alignment of *NbLIG1.1*- and *NbLIG1.2*-derived qPCR primers with the other *NbLIG1* isoform.

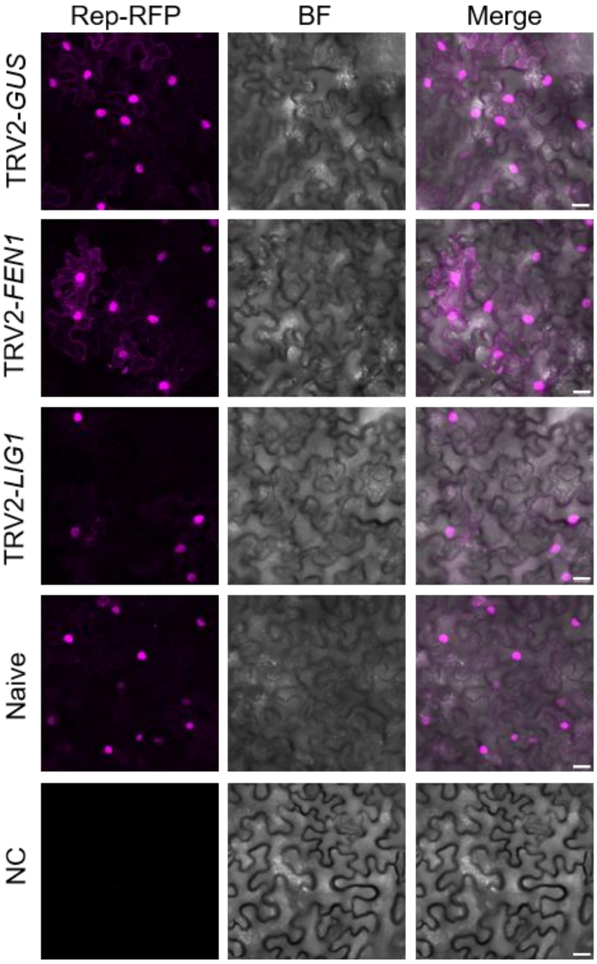


**Figure S17. *Agrobacterium tumefaciens*-mediated transient expression of Rep-RFP is not impaired in *FEN1*- and *LIG1*-silenced *N. benthamiana* plants.** *Agrobacterium* cells containing TRV1 and TRV2-FEN1and TRV2-LIG1 were mixed at a 1:1 ratio and inoculated into 2-week-old *N. benthamiana* seedlings. Plants were then agroinfiltrated with Rep-RFP at 14 days post-TRV inoculation. BF, brightfield. Merge, overlay of fluorescence and BF images. Scale bar: 20 μm. TRV2-GUS-inoculated and naïve samples served as positive controls, while NC represents the negative control (infiltrated with an empty vector). This experiment was performed in parallel with the experiments shown in Figure S5; therefore, the same negative control images (TRV2-GUS and naïve samples) are presented here. This experiment was repeated twice with similar results.

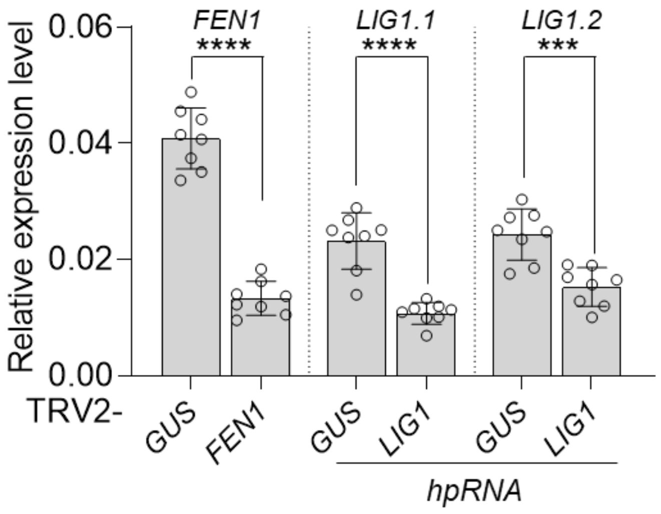


**Figure S18. Silencing efficiency in *FEN1*- and *LIG1*-silenced *N. benthamiana* plants in Figure 5.** qRT-PCR analysis of silencing efficiency in *FEN1*- and *LIG1*-silenced WT *N. benthamiana* plants shown in Figure 5 following agroinfiltration with the TYLCV infectious clone. *NbActin* served as an internal reference gene. Each dot represents an independent plant. Error bars indicate standard deviation. TRV2-GUS plants served as the control. Asterisks indicate statistically significant differences based on Student's t-test (****, P<0.0001; ***, P<0.001; **, P<0.01; *, P<0.05; not significant (ns), P>0.05). All experiments were repeated at least twice with similar results.

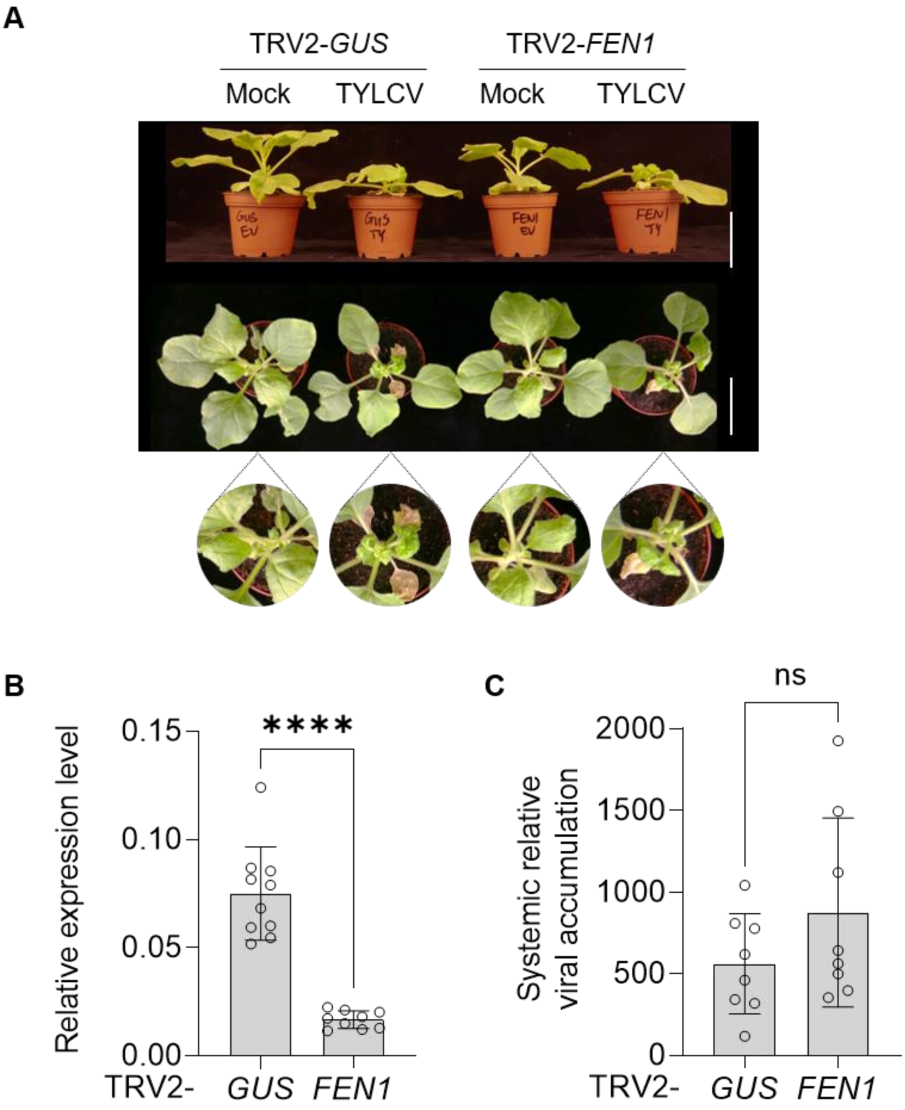


**Figure S19. FEN1 is not required for TYLCV systemic infection. A-C.** Symptoms (A, scale bar: 5 cm) in *FEN1*-silenced *N. benthamiana* plants upon TYLCV systemic infection. Silencing efficiency (B), measured by qRT-PCR using *NbActin* as an internal reference, and viral accumulation (C) measured by qPCR with 25S ribosomal DNA interspacer (ITS) as an internal reference. *Agrobacterium* cells containing the TYLCV infectious clone, TRV1, and TRV2 or TRV2-FEN1 were mixed at a 1:1:1 ratio and inoculated in 2-week-old *N. benthamiana* seedlings. Samples were harvested at 14 days post-inoculation. Plants infiltrated with TRV2-GUS were used as negative controls. These experiments were repeated three times with similar results. Each dot represents an independent plant. Error bars indicate standard deviation. TRV2-GUS plants served as the control. Asterisks indicate statistically significant differences based on Student's t-test (****, P<0.0001; ***, P<0.001; **, P<0.01; *, P<0.05; not significant (ns), P>0.05).

A

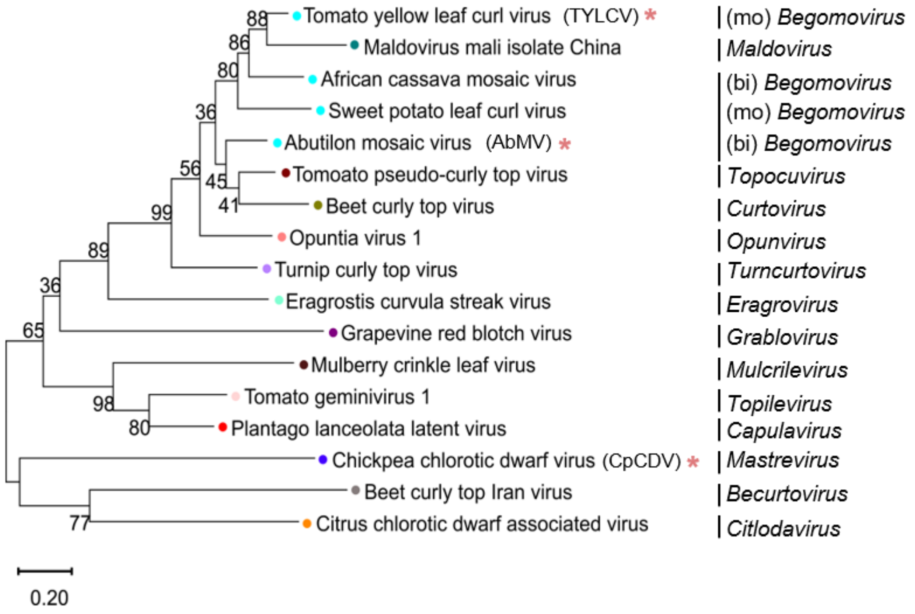


B

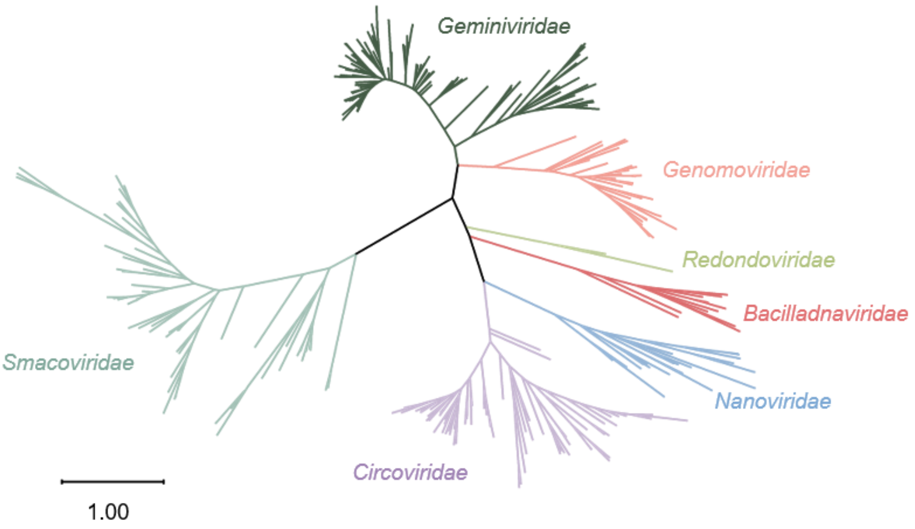


**Figure S20. Phylogenetic tree of geminiviruses and CRESS DNA viruses. A.** Phylogenetic tree based on Rep protein sequences of representative geminiviruses. The tree was built using the neighbor-joining method with 1,000 bootstrap replicates. Numbers at branches and the bottom indicate bootstrap values (%) and substitutions per site, respectively. Mono- and bipartite begomovirus were labelled as "mo" and "bi", respectively. Asterisks indicate the viruses used in this work. **B.** Maximum likelihood phylogenetic tree of multicellular eukaryote-infecting CRESS DNA viruses, based on the Rep protein sequence. The phylogenetic tree was constructed using the maximum likelihood method with 50 bootstrap replicates. Sequences were obtained from Kazlauskas et al. (2018) and supplemented with updated entries retrieved from the International Committee on Taxonomy of Viruses (ICTV) and the NCBI database. Viral families are color-coded and labelled in branches.

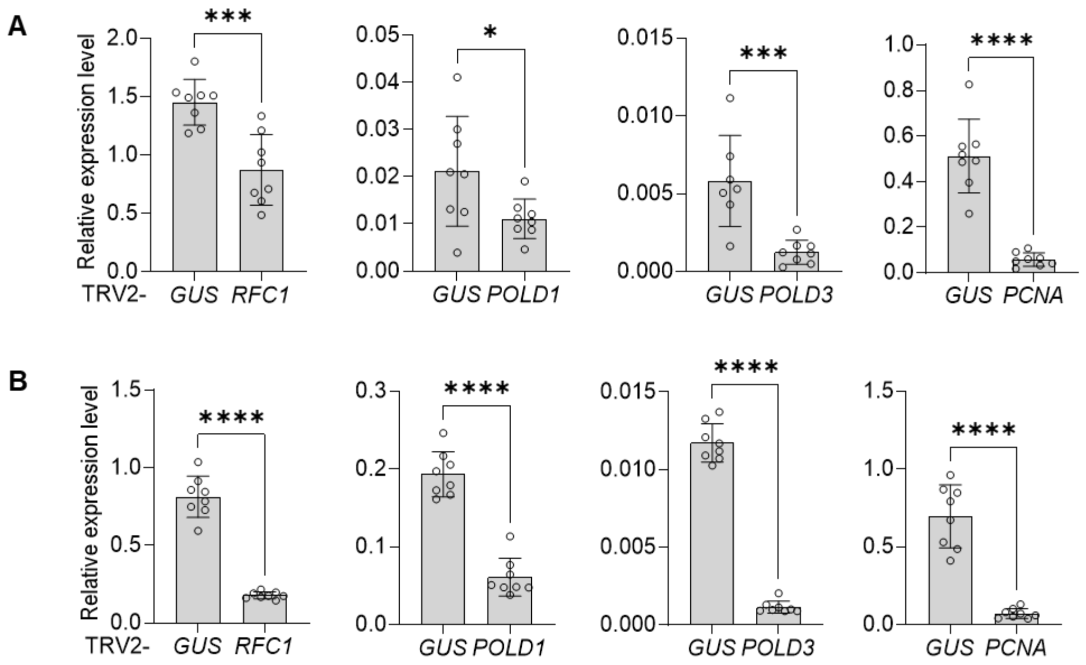


**Figure S21. Silencing efficiency in gene-silenced plants in Figure 6D and E. A-B.** qRT-PCR analysis of silencing efficiency in *RFC1*-, *POLD1*-, *POLD3*-, and *PCNA*-silenced WT *N. benthamiana* plants following CpCDV systemic infection (A) or agroinfiltration with PNYDV DNA-R (B). Panels A and B correspond to Figures 6D and 6E, respectively. *NbActin* served as an internal reference gene. These experiments were repeated twice with similar results. Each dot represents an independent plant. Error bars indicate standard deviation. TRV2-GUS plants served as the control. Asterisks indicate statistically significant differences based on Student's t-test (****, P<0.0001; ***, P<0.001; **, P<0.01; *, P<0.05; not significant (ns), P>0.05).

### SUPPLEMENTARY TABLES

**Supplementary Table 1. Datasets from TurboID-based PL experiments with the Rep proteins from TYLCV and AbMV in infected cells.**

**Supplementary Table 2. Datasets from TurboID-based PL experiments with the Rep proteins from TYLCV and AbMV in unifected cells.**

**Supplementary Table 3. Plasmids used in this study.**

**Supplementary Table 4. Primers used in this study**
